# Learning the Vector Coding of Egocentric Boundary Cells from Visual Data

**DOI:** 10.1101/2022.01.28.478267

**Authors:** Yanbo Lian, Simon Williams, Andrew S. Alexander, Michael E. Hasselmo, Anthony N. Burkitt

## Abstract

The use of spatial maps to navigate through the world requires a complex ongoing transformation of egocentric views of the environment into position within the allocentric map. Recent research has discovered neurons in retrosplenial cortex and other structures that could mediate the transformation from egocentric views to allocentric views. These egocentric boundary cells respond to the egocentric direction and distance of barriers relative to an animals point of view. This egocentric coding based on the visual features of barriers would seem to require complex dynamics of cortical interactions. However, computational models presented here show that egocentric boundary cells can be generated with a remarkably simple synaptic learning rule that forms a sparse representation of visual input as an animal explores the environment. Simulation of this simple sparse synaptic modification generates a population of egocentric boundary cells with distributions of direction and distance coding that strikingly resemble those observed within the retrosplenial cortex. This provides a framework for understanding the properties of neuronal populations in the retrosplenial cortex that may be essential for interfacing egocentric sensory information with allocentric spatial maps of the world formed by neurons in downstream areas including the grid cells in entorhinal cortex and place cells in the hippocampus.

## 1 Introduction

Animals can perform extremely complex spatial navigation tasks, but how the brain implements a navigational system to accomplish this remains largely unknown. In the past few decades, many functional cells that play an important role in spatial cognition have been discovered, including place cells (O’Keefe and Dostrovsky, 1971; O’Keefe, 1976), head direction cells (Taube et al., 1990a, b), grid cells (Hafting et al., 2005; Stensola et al., 2012), boundary cells (Solstad et al., 2008; Lever et al., 2009), and speed cells (Kropff et al., 2015; Hinman et al., 2016). All of these cells have been investigated in the allocentric reference frame that is viewpoint-invariant.

However, animals experience and learn about environmental features through exploration using sensory input that is in their egocentric reference frame. Recently, some egocentric spatial representations have been found in multiple brain areas such as lateral entorhinal cortex (Wang et al., 2018), postrhinal cortices (Gofman et al., 2019; LaChance et al., 2019), dorsal striatum (Hinman et al., 2019), and the retrosplenial cortex (RSC) (Wang et al., 2018; Alexander et al., 2020). In the studies by Hinman et al.(2019) and Alexander et al.(2020), a very interesting type of spatial cell, the egocentric boundary cell, was discovered. Similar to allocentric boundary cells (Solstad et al., 2008; Lever et al., 2009), egocentric boundary cells (EBCs) possess vectorial receptive fields sensitive to the bearing and distance of nearby walls or boundaries, but in the egocentric reference frame. For example, an EBC of a rat that responds whenever there is a wall at at particular distance on the left of the rat means that the response of the EBC not only depends on the location of the animal but also its running direction or head direction, i.e., the cell is tuned to a wall in the animal-centered reference frame.

Alexander et al.(2020) identified three categories of EBCs in the rat RSC: proximal EBC whose egocentric receptive field boundary is close to the animal, distal EBC whose egocentric receptive field boundary is further away from the animal, and inverse EBC that respond everywhere in the environment except when the animal is close to the boundary. Some examples of proximal, distal and inverse EBCs are shown in Figure 1. Furthermore, EBCs in this area display a considerable diversity in vector coding; namely the EBCs respond to egocentric boundaries at various orientations and distances. Somewhat surprisingly, there are also EBCs tuned to a wall that is behind the animal (see the bottom plot of Figure 1b for an example).

**Figure 1:**
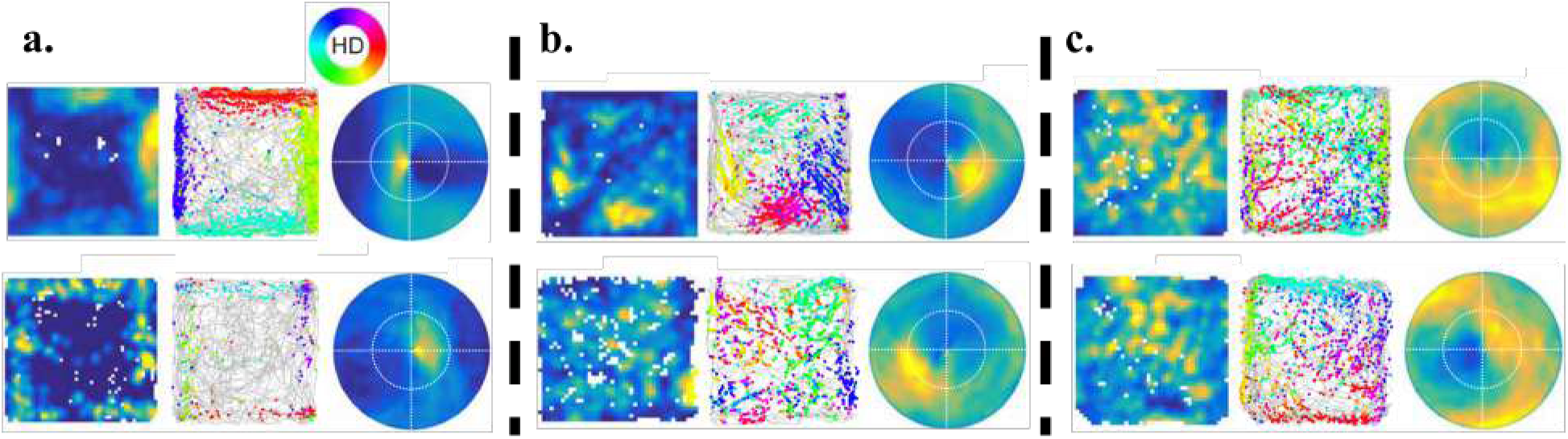
Six example EBCs from Alexander et al.(2020). The plots in the left column are the 2D spatial ratemaps, the middle column plots are trajectory plots showing firing locations and head directions (according to the circular color legend shown above **a**), and the right column plots are the receptive fields of the respective EBCs (front direction corresponds to top of page). **a)** Proximal EBCs whose receptive field is a wall close to the animal. The two example EBCs displayed here are selective to proximal walls of left and right, respectively. **b)** Distal EBCs whose receptive field is a wall further from the animal. The two example EBCs displayed here are selective to distal walls of rear-right and behind, respectively. **c)** Inverse EBCs that fire everywhere except when there is wall near the animal. The two example EBCs displayed here only stop firing when there are wall in front of and on the left of the animal, respectively.

Though there is increasing experimental evidence that suggests the importance of egocentric spatial cells, there have been few studies explaining how egocentric boundary cells are formed and whether they emerge from neural plasticity.

In this study, we show how EBCs can be generated through a learning process based upon sparse coding that uses visual information as the input. Furthermore, the learnt EBCs show a diversity of types, namely proximal, distal and inverse, and they fire for boundaries at different orientations and distances, similar to that observed in the experimental study of the vector coding for EBCs (Alexander et al., 2020). As Bicanski and Burgess (2020) pointed out in a recent review, the fact that some EBCs respond for boundaries behind the animal suggests that these cells do not solely rely on sensory input and appear to incorporate some mnemonic components. However, our model shows that by solely taking visual input, without any mnemonic component, some learnt EBCs respond to boundaries that are behind the animal and out of view.

These boundaries can, nevertheless, be inferred from distal visual cues, suggesting that the competition introduced by sparse coding drives different model cells to learn responses to boundaries at a wide range of different directions.

We next show that the model based on sparse coding that takes visual input while a simulated animal explores freely in a 2D environment can learn EBCs with diverse tuning properties and these learnt EBCs can generalize to novel environments.

## 2 Materials and Methods

### 2.1 The simulated environment, trajectory and visual input

#### 2.1.1 Environment

The simulated environment is programmed to match the experimental setup of Alexander et al.(2020) as closely as possible. It consists of a virtual walled arena 1.25 m by 1.25 m. One virtual wall is white and the other three are black. The floor is a lighter shade of grey with RGB values (0.4, 0.4, 0.4).

#### 2.1.2 Trajectory

The simulated trajectory is generated randomly using the parameters from Raudies and Hasselmo (2012). The simulated animal starts in the center of the arena facing north with the white wall to the right. This is used as the 0° bearing direction. The velocity of the animal is sampled from a Rayleigh distribution with mean 13 cm/s while enforcing a minimum speed of 5 cm/s.

The direction of motion is modelled by a random walk for the bearing, where the change in bearing at each time step is sampled from a zero mean normal distribution with standard deviation 340° per second and scaled to the length of the time step.

A complication for the simulation is how to deal with the walls. Following Raudies and Hasselmo (2012), we encode the following. If the simulated animal will approach within 2 cm of one of the walls on its next step, its velocity is adjusted to halfway between the current speed and the minimum speed (5 cm/s). Additionally we change the bearing by turning away from the wall by 90°.

#### 2.1.3 Visual input

The simulated environment and trajectory above are realised using the Panda3D game engine (`panda3d.org`), an open-source framework for creating virtual visual environments, usually for games. The visual input of the simulated animal is modelled using a camera with a 170° field of horizontal view to mimic the wide visual field of rat and a 110° field of vertical view. This input is used to generate a grayscale 8-bit image 170 × 110 pixels, which corresponds approximately to the visual acuity of the rat, namely 1 cycle per degree (Prusky et al., 2000). The camera is always facing front, meaning that the head direction is aligned with the movement direction for the simulated animal. The simulation is run at 30 frames per second until 40000 frames have been collected, which approximately corresponds to a running trajectory over a period of 1300 s (21 min, 40 s).

Model results shown in this paper are based on the visual input with 170° field of view (FOV) except Section 3.3 where different FOVs ( 60°, 90°, 120°, 150°, and 170°) are simulated to investigate how the width of FOV affects the distribution of learnt EBCs.

### 2.2 Learning egocentric boundary cells (EBCs)

#### 2.2.1 Non-negative sparse coding

Sparse coding (Olshausen and Field, 1996, 1997) was originally proposed to demonstrate that simple cells in the primary visual cortex (V1) encode visual input using an efficient representation. The essence of sparse coding is the assumption that neurons within a network can represent the sensory input using a linear combination of some relatively small set of basis features (Olshausen and Field, 1997). Along with its variant, non-negative sparse coding (Hoyer, 2003), the principle of sparse coding provides a compelling explanation for neurophysiological findings for many brain areas such as the retina, visual cortex, auditory cortex, olfactory cortex, somatosensory cortex and other areas (see Beyeler et al.(2019) for a review). Recently, sparse coding with non-negative constraint has been shown to provide an account for learning of the spatial and temporal properties of hippocampal place cells within the entorhinal-hippocampal network (Lian and Burkitt, 2021, 2022). In this study, non-negative sparse coding is used to learn the receptive field properties of EBCs found in the RSC.

#### 2.2.2 Model structures

As the simulated animal runs freely in the 2D environment, an image representing what the animal sees is generated at every location. This image is used as the visual stimulus to the simulated animal. To explore where in the visual processing chain EBCs arise we investigate two models: (i) Raw Visual (RV) model, a control model that uses the raw visual data (model structure shown in Figure 2a), and (ii) V1-RSC model, a more biological model that uses the processed data corresponding to processing in the early visual system and processing in the V1 before projecting to the RSC (model structure shown in Figure 2b).

**Figure 2:**
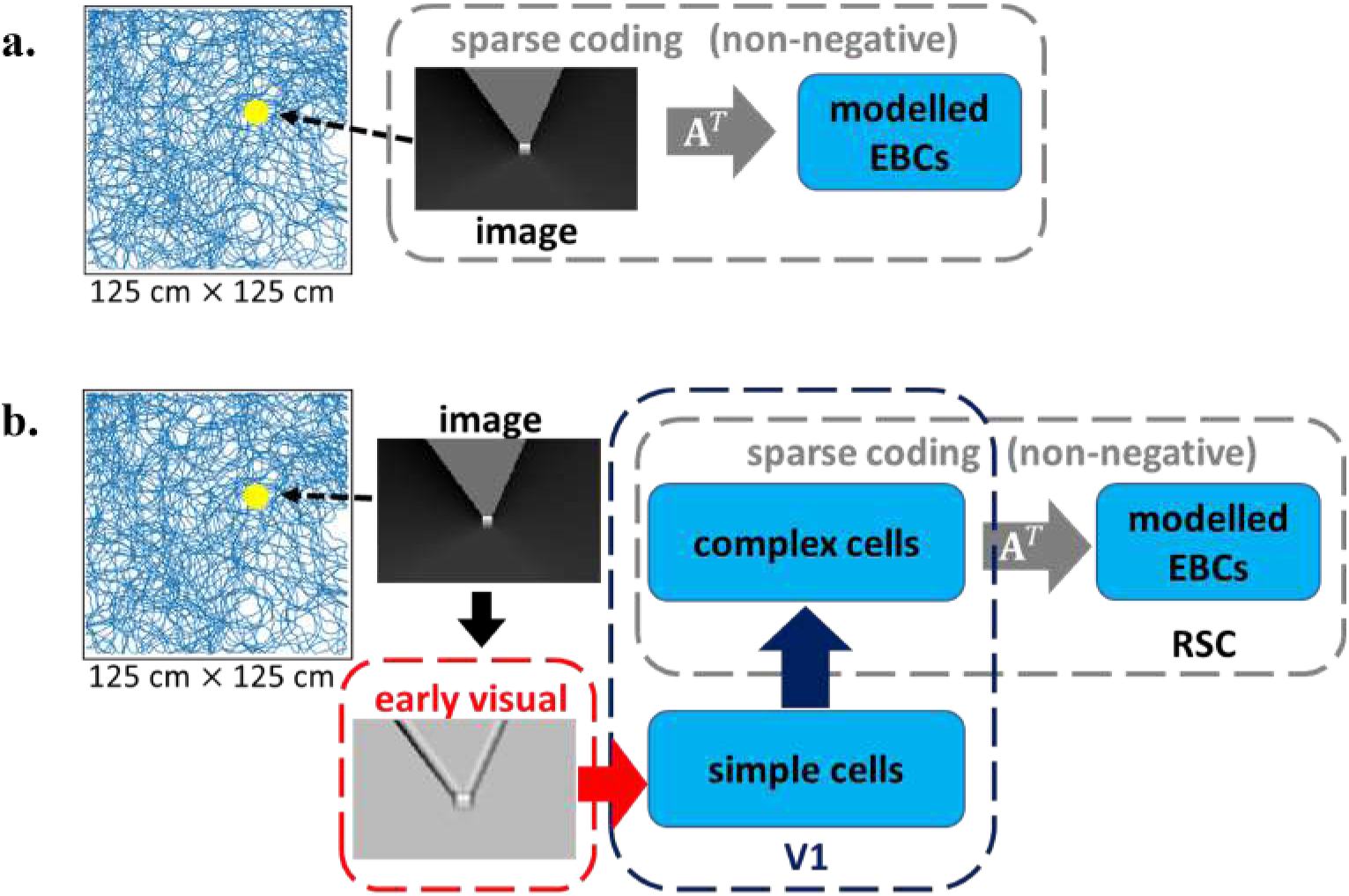
Structures of Raw Visual (RV) model and V1-RSC model. The simulated animal runs freely in the 1.25 m×1.25 m simulated environment. The simulated visual scene the animal sees at different locations is the visual stimulus to the simulated animal. a) RV model: the raw visual input is directly used as the input to a network that implements non-negative sparse coding. b) V1-RSC model: the raw visual input is preprocessed by the early visual system and then projected to V1 that involves simple cell and then complex cell processing; complex cells in V1 then project to modelled EBCs in RSC and a V1-RSC network is implemented based on non-negative sparse coding (described in Equations 2 & 3).

The learning principle used in both the RV and V1-RSC models is non-negative sparse coding. Given that the RV model is designed as a control model to investigate whether raw visual input can give rise to EBCs, while the V1-RSC model is a more biological model that incorporates visual processing in the early visual systems and V1, we use slightly different implementations of non-negative sparse coding. Specifically, the RV model uses a built-in function from the SciKit-Learn python package (Pedregosa et al., 2011) while the V1-RSC model uses the implementation from our previous work (Lian and Burkitt, 2021).

#### 2.2.3 Raw Visual model: using the raw visual data

In the RV model, the raw visual data is used as the input to the model, which is a 40000×18700 matrix. This contains the raw visual data (170 × 110) flattened for all 40000 time steps. One sample of raw visual input is displayed as the embedded ‘image’ in Figure 2a. Non-negative sparse coding of this model is implemented by applying non-negative matrix factorisation (Lee and Seung, 1999) with sparsity constraints using the built-in function from the SciKit-Learn python package (Pedregosa et al., 2011). 100 dictionary elements are generated, which we identify with the model neuron responses used in the V1-RSC model. Since the simulated animal only has access to the visual data as it is running, training on the 40000 × 18700 input dataset is only performed for a single iteration to simulate the early stages of receptive field generation.

#### 2.2.4 V1-RSC model: using a more biological model from V1 to RSC

**Early visual processing:** Processing in the early visual system describes the visual processing of the retinal ganglion cells (RGCs). In this study, this is done using divisively normalised difference-of-Gaussian filters that mimic the receptive fields of RGCs in the early visual system (Tadmor and Tolhurst, 2000; Ratliff et al., 2010). For any input image, the filtered image *I* at point (*x, y*) is given by

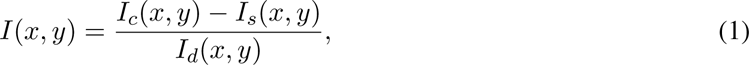

where *I_c_*, *I_s_*, and *I_d_* are the response of the input image filtered by three unit-normalised Gaussian filters: center filter (*G_c_*), surround filter (*G_s_*), and divisive normalisation filter (*G_d_*). *G_c_* `−`*G_s_* implements the typical difference-of-Gaussian filter that characterises the center-surround receptive field of retinal ganglion cells and *G_d_* describes the local adaptation of RGCs (Troy et al., 1993). The receptive field size of RGCs is set to 9 × 9. The standard deviations of *G_c_*, *G_s_* and *G_d_* are set to 1, 1.5 and 1.5, respectively (Borghuis et al., 2008). RGCs are located at each pixel point of the input image except these points that are within 4 pixels of the edges of the input image. For a given input image with size 170 × 110, the processed image after the early visual system has size 162 × 102. One sample of raw visual input and its corresponding processed input by the early visual system are displayed as the embedded ‘image’ and ‘early visual’ in Figure 2b.

**V1 processing:** Next, visual information processed by the early visual system projects to V1 and is further processed by simple cells and complex cells in V1 (Lian et al., 2019, 2021). The receptive field of a simple or complex cell is characterised by a 13 × 13 image. Simple cells are described as Gabor filters with orientations spanning from 0° to 150° with step size of 30°, spatial frequencies spanning from 0.1 to 0.2 cycles per pixel with step size of 0.025, and spatial phases of 0°, 90°, 180° and 270°. In addition, a complex cell receives input from 4 simple cells that have the same orientation and spatial frequency but different spatial phases (Movshon et al., 1978a, b; Carandini, 2006). Therefore, at each location of a receptive field, there are 6 × 5 × 4 = 120 simple cells and 6 × 5 = 30 complex cells. As the receptive field only covers a small part of the visual field, the same simple cells and complex cells are repeated after every 5 pixels. Given that an input image from the early visual system has size 162 × 102 and the size of a receptive field is 13 × 13, there are 27 × 20 = 540 locations that have simple cells and complex cells. Overall, there are 120 × 540 = 64800 simple cells and 30 × 540 = 16200 complex cells in total. For a given visual stimulus with size 170 × 110, complex cell responses can be represented by a 16200 × 1 vector. After the vision processing in V1, complex cell responses in V1 project to the RSC.

**Model dynamics:** Similar to our previous work (Lian and Burkitt, 2021, 2022), we implement the model via a locally competitive algorithm (Rozell et al., 2008) that efficiently solves sparse coding as follows:

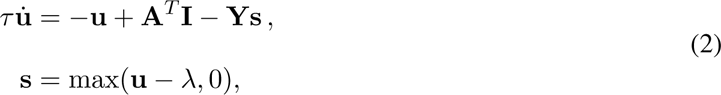

and

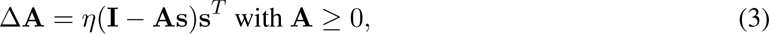

where **I** is the input from V1 (i.e., complex cells responses), **s** represent the response (firing rate) of the model neurons in the RSC, **u** can be interpreted as the corresponding membrane potential, **A** is the matrix containing basis vectors and can be interpreted as the connection weights between complex cells in V1 and model neurons in the RSC, 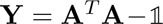 and can be interpreted as the recurrent connection between model neurons in the RSC, 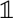 is the identity matrix, *τ* is the time constant of the model neurons in the RSC, *λ* is the positive sparsity constant that controls the threshold of firing, and *η* is the learning rate. Each column of **A** is normalised to have length 1. Non-negativity of both **s** and **A** in Equations 2 & 3 is incorporated to implement non-negative sparse coding. Additional details about the above implementation of non-negative sparse coding can be found in Lian and Burkitt (2021).

**Training:** For the implementation of this model, there are 100 model RSC neurons and the parameters are given below. For the model dynamics and learning rule described in Equations 2 & 3, *τ* is 10 ms, *λ* is 0, and the time step of implementing the model dynamics is 0.5 ms. The simulated visual input of the simulated trajectory that contains 40000 positions is used to train the model. Since the simulated trajectory is updated after every 30 ms, at each position of the trajectory, there are 60 iterations of computing the model response using Equation 2. After these 60 iterations, the learning rule in Equation 3 is applied such that connection **A** is updated. The animal then moves to the next position of the simulated trajectory. The learning rate *η* is set to 0.3 for the first 75% of the simulated trajectory and 0.03 for the final 25% of the simulated trajectory.

Note that the model with *λ* = 0 implements non-negative matrix factorisation (Lee and Seung, 1999), which is a special variant of non-negative sparse coding. However, when *λ* is set to a positive value such as 0.1, the learnt EBCs display similar features, except that the neural response is sparser.

### 2.3 Collecting model data

After the RV model and V1-RSC model finish learning using simulated visual input sampled along the simulated trajectory, a testing trajectory with simulated visual input is used to collect model responses for further data analysis. The experimental trajectory of real rats from Alexander et al.(2020) is used as the testing trajectory and it contains movement direction as well as head direction. In addition, for the experimental trajectory, head direction is not necessarily identical to movement direction because the animal is not head-fixed in the experiment. Simulated visual input from the experimental trajectory is generated using the same approach described above, except that the camera is not facing front but aligned with the head direction from the experimental data. Both models are rate-based and thus the model responses are then transformed into spikes using a Poisson spike generator with a maximum firing rate 30 Hz for the whole modelled population.

Results displayed in the main text are generated using model data collected from an experimental trajectory that has different movement and head directions. However, results of model data collected from a simulated trajectory where head direction is aligned with movement direction are also given in Supplementary Materials.

### 2.4 Experimental methods

An electrophysiological dataset collected from the RSC of male rats performing random foraging in a 1.25 m×1.25 m arena was used from published prior work (Alexander et al., 2020) to make comparisons between model and experiment data of EBCs. For additional details relating to experimental data acquisition see Alexander et al.(2020). In addition, the data analysis techniques from this experimental paper were used to analyze the data from the simulations.

### 2.5 Data analysis

#### 2.5.1 Two-dimensional (2D) spatial ratemaps and spatial stability

The analysis of the neural activity in the simulation used the same techniques that were used to analyze published experimental data from the RSC (Alexander et al., 2020). Animal or simulation positional occupancy within an open field was discretized into 3 cm×3 cm spatial bins. For each model neuron, the raw firing rate for each spatial bin was calculated by dividing the number of spikes that occurred in a given bin by the amount of time the animal occupied that bin. Note that spiking in the model was generated by a Poisson spike generator. Raw firing ratemaps were smoothed with a 2D Gaussian kernel spanning 3 cm to generate final ratemaps for visualization.

#### 2.5.2 Construction of egocentric boundary ratemaps

The analysis of egocentric boundary ratemaps (EBR) used the same techniques used for published experimental data (Alexander et al., 2020). EBRs were computed in a manner similar to 2D spatial ratemaps, but referenced relative to the animal rather than the spatial environment. The position of the boundaries relative to the animal was calculated for each position sample (i.e., frame). For each frame, we found the distance, in 2.5 cm bins, between arena boundaries and angles radiating from 0° to 360° in 3° bins relative to the animal’s position. Angular bins were referenced to the head direction of the animal such that 0°*/*360° was always directly in front of the animal, 90° to its left, 180° directly behind it, and 270° to its right. Intersections between each angle and environmental boundaries were only considered if the distance to intersection was less than or equal to half the length to the most distant possible boundary (in most cases this threshold was set at 62.5 cm or half the width of the arena to avoid ambiguity about the influence of opposite walls). In any frame, the animal occupied a specific distance and angle relative to multiple locations along the arena boundaries, and accordingly, for each frame, the presence of multiple boundary locations were added to multiple 3° × 2.5 cm bins in the egocentric boundary occupancy map. The same process was completed with the locations of individual spikes from each model neuron, and an EBR was constructed by dividing the number of spikes in each 3° × 2.5 cm bin by the amount of time that bin was occupied in seconds. Smoothed EBRs were calculated by convolving each raw EBR with a 2D Gaussian kernel (5 bin width, 5 bin standard deviation).

#### 2.5.3 Identification of neurons with egocentric boundary vector tuning

The identification of model neurons with significant egocentric boundary vector sensitivity used the same criteria for identification of real neurons showing this response (Alexander et al., 2020). The mean resultant, 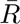 of the cell’s egocentric boundary directional firing, collapsed across distance to the boundary, was calculated as

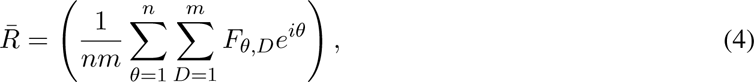

where *θ* is the orientation relative to the rat, *D* is the distance from the rat, *F_θ,D_*is the firing rate in a given orientation-by-distance bin, *n* is the number of orientation bins, and *m* is the number of distance bins. The mean resultant length (MRL), 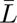, is defined as the absolute value of the mean resultant and characterized the strength of egocentric bearing tuning to environment boundaries. The preferred orientation of the egocentric boundary ratemap was calculated as the mean resultant angle (MRA), 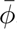,

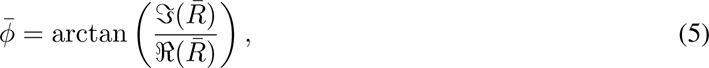

where 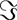 and 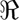 are the real and imaginary parts of their arguments respectively.

The preferred distance was estimated by fitting a Weibull distribution to the firing rate vector corresponding to the MRA and finding the distance bin with the maximum firing rate. The MRL, MRA, and preferred distance were calculated for each model neuron for the two halves of the experimental session independently.

A model neuron was characterized as having egocentric boundary vector tuning (i.e., an EBC) if it reached the following criteria: 1) the MRL from both session halves were greater than the 99th percentile of the randomized distribution taken from Alexander et al.(2020) (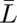 *>* 0.14), 2) the absolute circular distance in preferred angle between the 1st and 2nd halves of the baseline session was less than 45°, and 3) the change in preferred distance for both the 1st and 2nd halves relative to the full session was less than 50%. To refine our estimate of the preferred orientation and preferred distance of each model neuron we calculated the center of mass (COM) of the receptive field defined after thresholding the entire EBR at 75% of the peak firing and finding the largest continuous contour (‘contour’ in Matlab). We repeated the same process for the inverse EBR for all cells to identify both an excitatory and inhibitory receptive field and corresponding preferred orientation and distance for each model neuron.

#### 2.5.4 Von Mises mixture models

Distribution of preferred orientation estimates was modeled as mixtures of Von Mises distributions using orders from 1 to 5 (“fitmvmdist” found at https://github.com/chrschy/mvmdist). Optimal models were identified as the simplest model increasing model fit by 10% over the one-component model. Theta of each Von Mises component is reported, and a distribution function of the optimal model was generated to visualize mixture model fit.

## 3 Results

### 3.1 Learnt EBCs are similar to those found in the experimental study

#### 3.1.1 Results using Raw Visual model

100 dictionary elements (model cells) of the RV model were trained on a simulated trajectory and then tested on the experimental trajectory as described in Section 2.3. 38% of these model cells possessed significant and reliable sensitivity to the egocentric bearing and distance to environmental boundaries. A similar but lightly larger percentage was observed when these model cells were tested on the simulated trajectory (41%). Figure 3 shows six examples of learnt cells that are proximal, distal and inverse EBCs. Plots of the full set of 100 RV model cells tested using experimental and simulated animal trajectories are given in the Materials A.1 & A.2.

**Figure 3:**
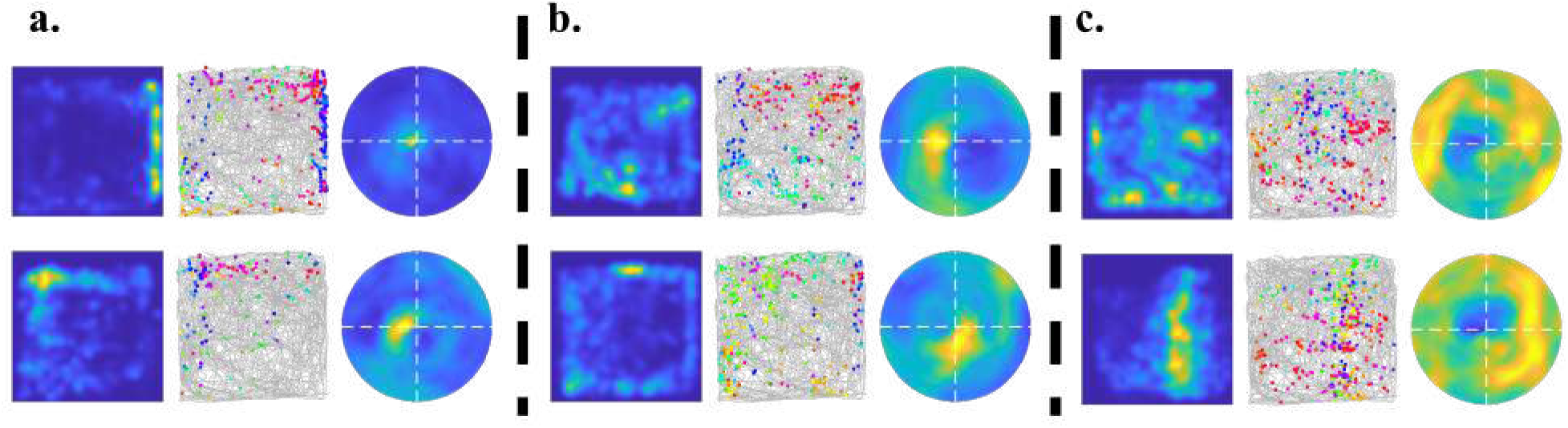
Examples of learnt EBCs recovered using experimental trajectory: Raw Visual model. Similar to Figure 1, each row with three images shows the spatial ratemap, firing plot with head directions and egocentric ratemap. **a)** Proximal EBCs, **b)** Distal EBCs, and **c)** Inverse EBCs with different preferences of egocentric orientation.

#### 3.1.2 Results using V1-RSC model

100 model cells of the V1-RSC model were also trained using on a simulated trajectory and then tested on the experimental trajectory, as described in Section 2.3. Of these cells, 85% possessed significant egocentric boundary vector sensitivity when tested on the real animal trajectory and a similar percentage was observed on the simulated trajectory (90%). Twelve examples showing the activity of cells with learned EBC receptive fields on the experimental trajectory are displayed in Figure 4. The four sets of plots in Figure 4a depict representative examples of proximal EBCs with different preferences for egocentric orientation, and the four sets of plots in Figure 4b show representative examples of distal EBCs, also showing different preferences for egocentric orientation. The four sets of plots in Figure 4c show examples of learned inverse EBCs. Each row consists of EBCs with similar orientations. These examples illustrate that they code for different orientations and distances in the animal-centered framework. Plots of the full set of 100 V1-RSC model cells generated using experimental and simulated animal trajectories are given in the Supplementary Materials A.3 & A.4.

**Figure 4:**
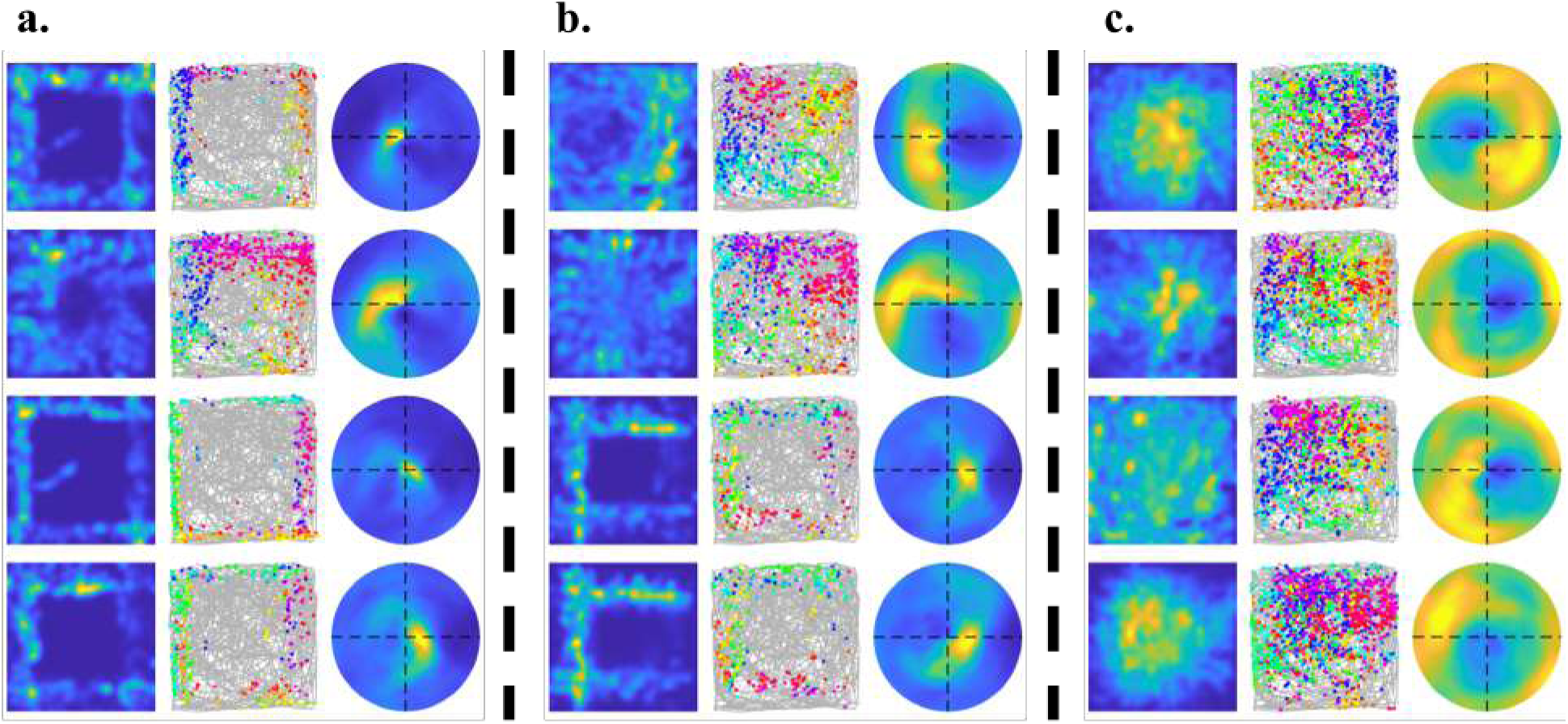
Examples of learnt EBCs recovered using experimental trajectory: V1-RSC model. Similar to Figure 1, each row with three images shows the spatial ratemap, firing plot with head directions and egocentric ratemap. **a)** Proximal EBCs, **b)** Distal EBCs, and **c)** Inverse EBCs with different preferences of egocentric orientation.

These result show that, after training, the learnt RSC cells exhibit diverse egocentric tuning similar to that observed in experimental data (Alexander et al., 2020), including the three different types identified experimentally: proximal, distal and inverse. The results likewise show that the cells are activated by walls at different orientations in the egocentric framework. In other words, this model learns diverse egocentric vector coding; namely the learnt cells code for boundaries at different orientations and distances.

#### 3.1.3 Population statistics of EBC orientation and distance

The EBCs that are learnt using RV model and V1-RSC model, illustrated in Figure s3 & 4, show considerable similarity to those found in experimental studies (Alexander et al., 2020). After the model is trained on simulated visual data sampled from a virtual environment with a simulated trajectory, model responses are collected with both experimental trajectory (where head direction is not necessarily aligned with moving direction) and simulated trajectory (where head direction is the same as moving direction), see “Collecting model data” of “Materials and Methods” for details. Then the egocentric tuning properties of all the model cells are investigated using the technique in “Data analysis” of “Materials and Methods”.

A summary of percentages of cells that are classified as EBCs for both experimental and model data is displayed in Table 1. Alexander et al.(2020) reported 24.1% (n=134/555) EBCs in the experimental data. RV model has 41% (n=41/100) and 38% (n=38/100) EBCs recovered by simulated trajectory and experimental trajectory, respectively. V1-RSC model has 90% (n=90/100) and 85% (n=85/100) EBCs recovered by simulated trajectory and experimental trajectory, respectively. Above all, our proposed model is successful in learning EBCs from visual input.

**Table 1:**
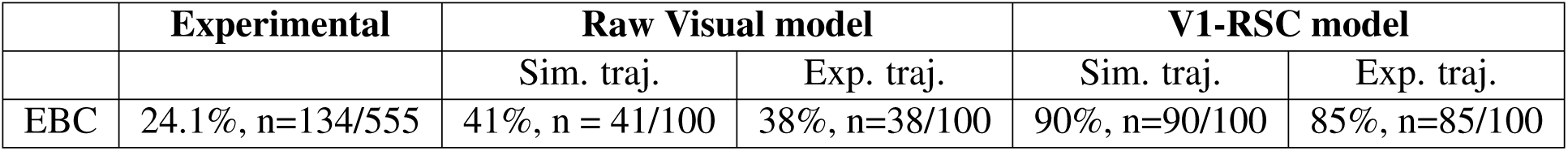
Percentages of EBCs of experimental and model data.

The extent of the similarity between experimental and model data is shown in Figure 5, which demonstrates that both RV and V1-RSC models generate EBCs whose characteristics resemble experimentally observed data on a population level. Thus, visual input alone may give rise to EBC-like receptive fields. The vector coding of an EBC indicates the coding of orientation and distance. Experimental data (left of Figure 5) shows that EBCs in the RSC have a lateral preference for orientation and a wide range of distance tuning. Learnt EBCs of both the RV model and V1-RSC model have qualitatively similar distributions to the experimental data of both preferred bearing and distance. That said, the distribution of preferred orientations and distances in the experimental dataset significantly differed from EBCs in the V1-RSC (Kuiper test for differences in preferred orientation; k = 3443; p = 0.002; Wilcoxon ranksum test for differences in preferred distance; p = 0.03) but not the RV model (Kuiper test for preferred orientation; k = 1644; p = 0.05; Wilcoxon ranksum test for preferred distance; p = 0.49). These differences partly arise from 1) an overall lack of V1-RSC EBCs with preferred egocentric orientations in front of or behind the animal and 2) a more uniform distribution of preferred distances with lower concentration in the proximal range for V1-RSC model EBCs.

**Figure 5:**
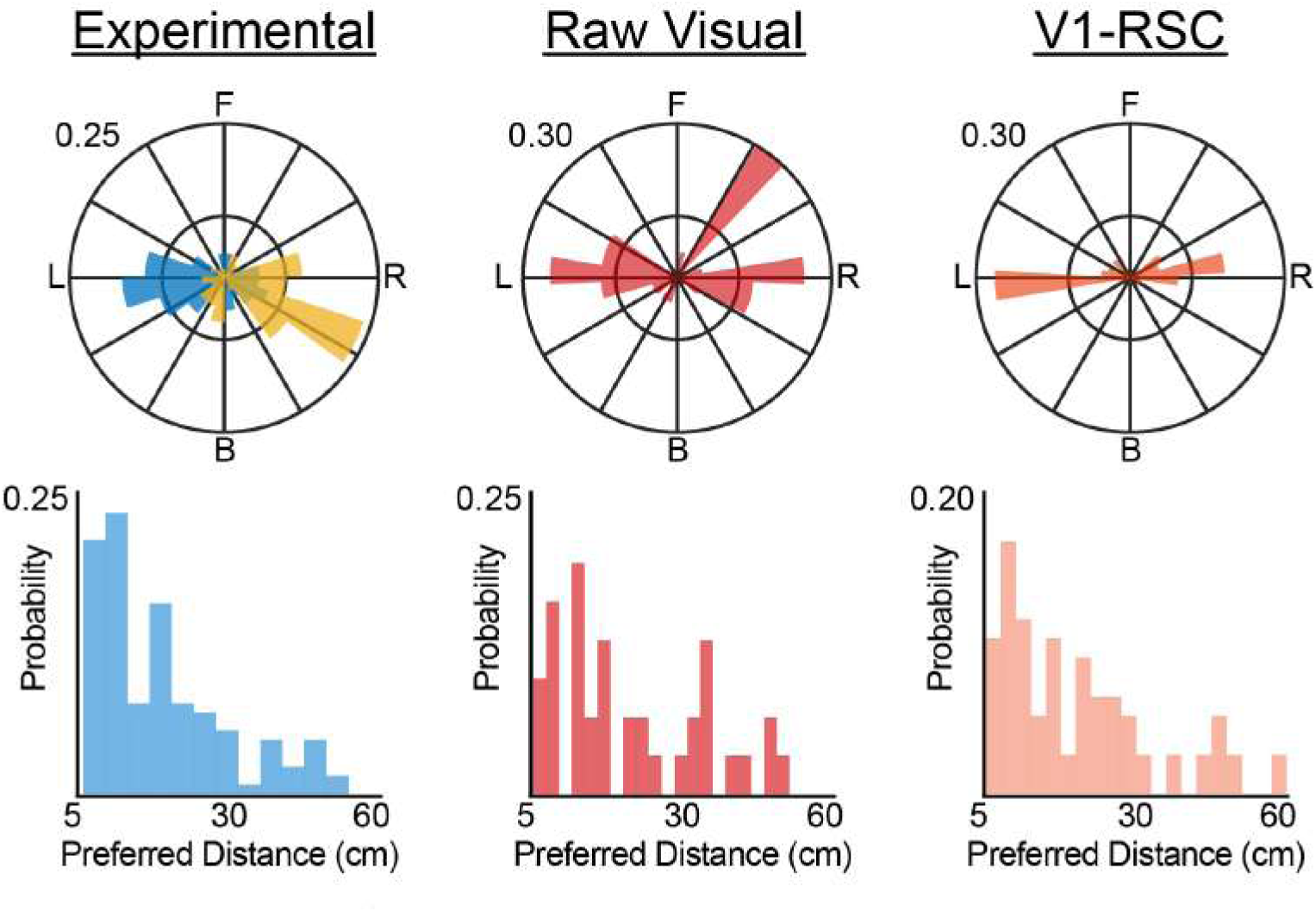
Population statistics of experimental and model data. Distributions of orientation (top row) and distance (bottom row) in the Raw Visual model (middle column) and V1-RSC model (right column) resemble experimental distributions observed in RSC (Alexander et al., 2020) (left column; blue and yellow histograms correspond to real neurons recorded in the right and left hemispheres, respectively). Model data in this Figure is collected using experimental trajectory.

Different visual inputs imply different spatial information about the animals’ position, so salient visual features may correlate with spatial tuning properties of neurons. By solely taking visual input, the model based on sparse coding promotes diverse tuning properties (different types of EBCs and diverse population responses) because of the inherent competition of the model. Difference between experimental and model data is discussed further in the Discussion Section 4.2 & 4.4.

### 3.2 Learnt EBCs generalize to novel environments

EBCs are experimentally observed to exhibit consistent tuning preferences across environments of different shapes or sizes (LaChance et al., 2019; Alexander et al., 2020). We next examined whether learnt EBCs of the two models exhibited similar characteristics. To do so, we exposed model units that were trained on the baseline (1.25m^2^) session to both a circular and expanded (2m^2^) novel environments.

We observed many learnt units that continued to exhibit egocentric receptive fields across environments (Figure 6a-e). However, there were notable differences in the preferred egocentric bearing and distances of the receptive fields of individual units as well as the generalizability of tuning across environments between the unprocessed (RV) and feature processed (V1-RSC) models. The RV model tended to have greater turnover of units with EBC-like properties between the baseline, circle, and expanded arenas while the population of EBCs in the V1-RSC model overlapped substantially between environments (e.g., only 1 RSC-V1 unit was an EBC solely in the baseline session; Figure 6f). Interestingly, both models exhibited more robust egocentric bearing tuning in circular when compared to square environments (Figure 6g; Kruskall-Wallis test w/ post-hoc Tukey-Kramer; RV *χ*^2^ = 42.3; V1-RSC *χ*^2^ = 63.5; both p *<*0.001). Consistent with this observation, RV model units were more likely to exhibit EBC-like tuning in circular environments (Figure 6b,f) while V1-RSC model units showed no preference for environment shape (Figure 6f).

**Figure 6:**
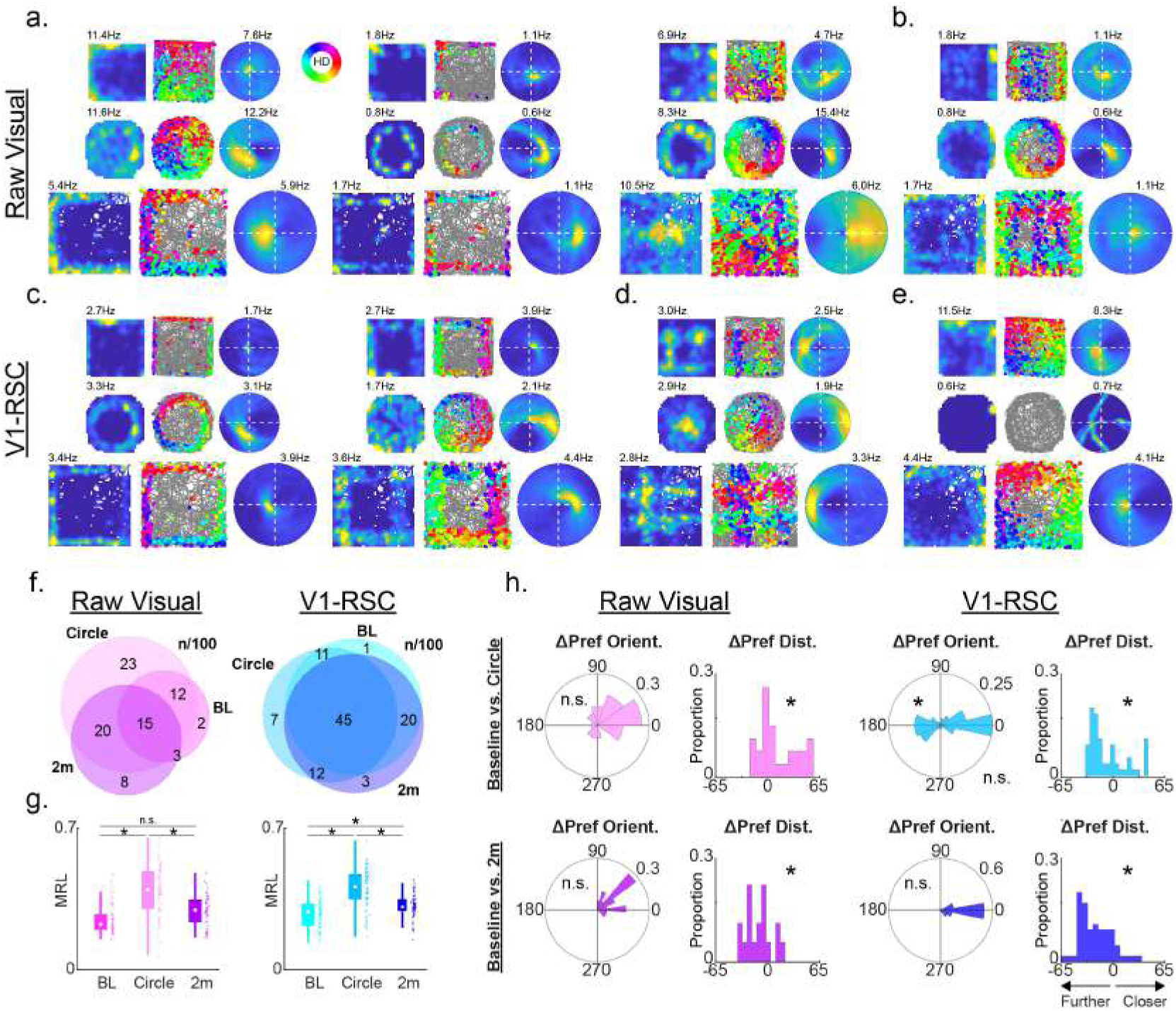
Model EBCs exhibit mostly consistent tuning when the environment is manipulated. **a)** 3 examples of EBCs in the Raw Visual (RV) model across baseline (1.25m^2^), circular, and expanded (2m^2^) environments. Left plots, firing ratemap as a function of position of the agent. Middle plots, trajectory plot showing agent path in gray and position at time of spiking as colored circles. Color indicates heading at the time of the spike as indicated in the legend. Right plots, egocentric boundary ratemap. **b)** RV unit with EBC coding in circular but not square environments. **c)** 2 examples of EBCs in the V1-RSC model across baseline (1.25m^2^), circular, and expanded (2m^2^) environments. Plots as in **a**. **d)** V1-RSC unit that has contralateral orientation tuning between square and circular environments. **e)** V1-RSC unit that loses an EBC receptive field when moving from square to circular environments. **f)** Venn diagrams for RV (left) and V1-RSC (right) EBCs across all simulated arenas. Overlaps indicate units with EBC tuning in multiple arenas. Numbers indicate total count out of 100 simulated units. BL, baseline (1.25m^2^); Circle, circular; 2m, expansion (2m^2^). **g)** Scatter plots of mean resultant length (MRL) for detected EBCs in each environment. Abbreviations as in **f**. **h)** Changes to preferred orientation and distance in RV and V1-RSC model EBC units between baseline and manipulation sessions. Rows are ‘baseline versus circle’ (top) or ‘baseline versus 2 meter’ (bottom) comparisons. Left four plots, RV model with polar plots depicting change to preferred orientation (Δ Pref Orient. = PO_baseline_ - PO_manip_) and histograms depicting change to preferred distance (Δ Pref Dist. = PD_baseline_ - PD_manip_). Radial and y-axes are the proportion of units with EBC-like tuning in both conditions. Negative values on the right histograms indicate receptive fields moving farther from the animal, vice versa for positive values. Right four plots, same as left plots but for the V1-RSC model.

The RV and V1-RSC models also diverged when examining the properties of egocentric boundary tuning curves across environments. While there were fewer preserved EBC units in the RV model across sessions, those that did maintain EBC-like tuning tended to have the similar preferred orientations between baseline, circular, and expanded arenas (Figure 6h, left column; Kuiper test for different preferred orientations; k _circle_ = 270; k_2m_ = 144; both p = 1). In contrast, V1-RSC units had significant differences at the population level in preferred orientations between the circular environment and baseline session (Figure 6h, top right; Kuiper test; k_circle_ = 1334; p = 0.001). This likely arose from subsets of V1-RSC units that exhibited movement of their preferred egocentric bearing to the contralateral side of the agent between arenas (Figure 6d,h). V1-RSC units were extremely reliable in their preferred orientation within both sized square arenas, indicating that the egocentric receptive fields in this model were highly sensitive to environmental geometry (Figure 6h, bottom right; Kuiper test; k_2m_ = 1105; p = 1). In fact, small numbers of V1-RSC units with EBC-like tuning in square environments exhibited a complete disruption of egocentric receptive fields in circular environments consistent with experimental observations (Figure 6e; A.S. Alexander, unpublished).

Larger alterations to EBC receptive fields across environments were observed for the distance component in both models. Many units exhibited drastic changes to their preferred egocentric distance with a bias towards a shift further from the animal (Figure 6h; Δ Pref Dist. = PD_baseline_ - PD_manip_; Signed rank test for 0 median differences; all conditions and models p*<*0.05). This observation was especially apparent in the V1-RSC model and, in particular, in the arena expansion manipulation (Figure 6h, bottom right). In the 2m ^2^ environment, shifts in preferred distances that moved receptive fields further away from the animal could indicate that subsets of EBCs anchored their activity to the center of the environment rather than boundaries, as reported in postrhinal cortices (Figure 6d; LaChance et al.2019). These simulations indicate that, in a manner consistent with experimentally observed EBCs, most model-derived units exhibit consistent EBC-like tuning between environments of different shapes and sizes.

### 3.3 The width of visual field affects the orientation distribution of learnt EBCs

The preferred egocentric bearings of EBCs from both experimental data and model simulations are concentrated at lateral angles (Figure 5) and overlap significantly with the facing direction of the eyes. Thus, it is possible that the distribution of EBC-preferred bearings reflects the visual field of the animal. We next examined model EBC receptive field properties in simulations of agents possessing varying fields of view (FOV, Figure 7). Consistent with this hypothesis, the distribution of preferred bearings is primarily forward facing in simulations with convergent FOVs and spread in more lateral orientations as the visual field approaches a more naturalistic width. Indeed, at a 170° width field of view, the distribution of preferred orientations becomes bimodal in both models with mean angular preferences of each mode falling near 0/360° and 180° as observed in experimental data (Figure 5). Accordingly, the combination of visual sparse coding and physical constraints on animal visual fields may define core properties of EBC receptive fields and enable the prediction of preferred bearings in other species.

**Figure 7:**
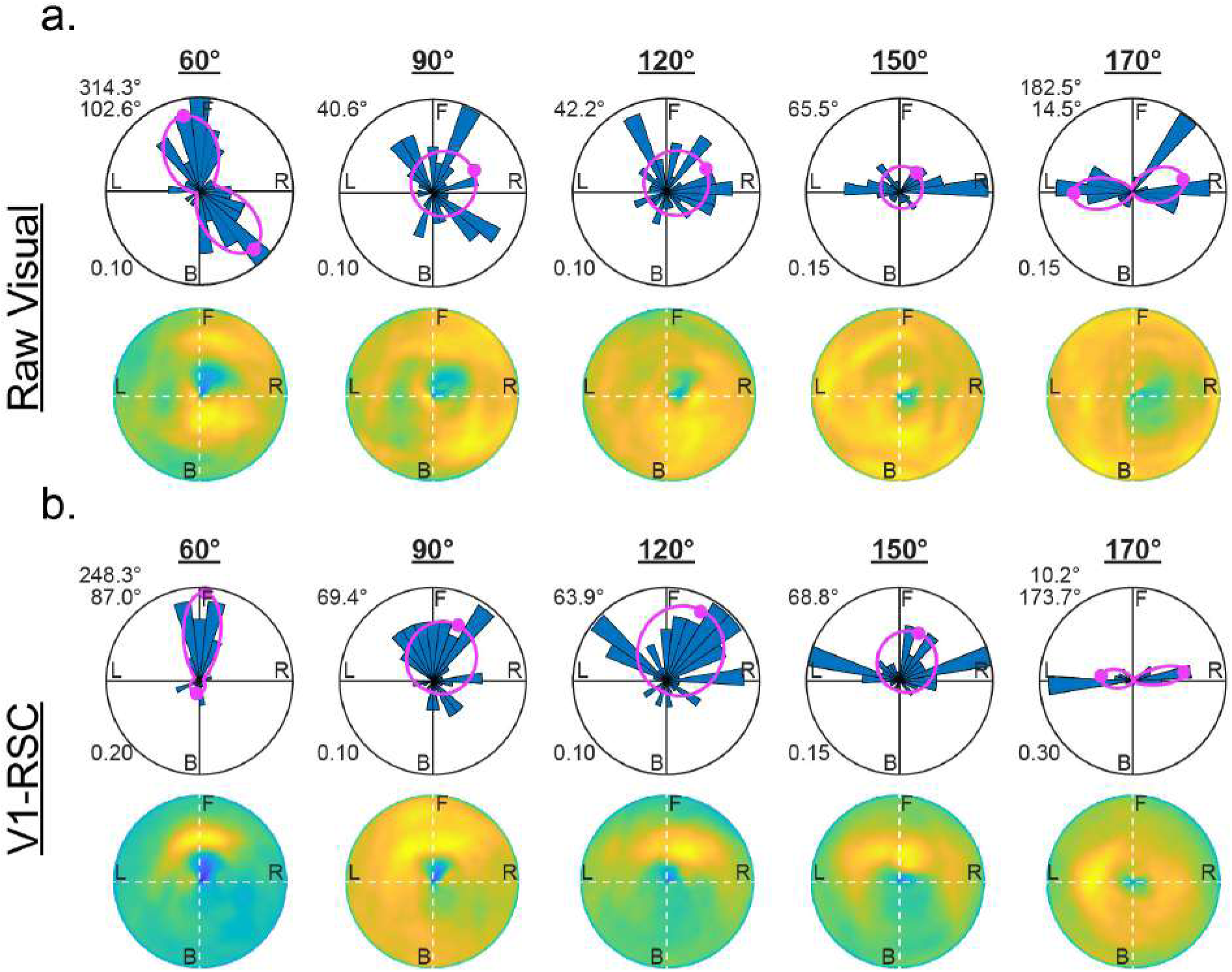
EBC preferred bearings as a function of field of view. **a)** Top row, distribution of preferred egocentric bearings for EBCs in the Raw Visual model as a function of width of field of view. Preferred bearings move from forward to lateral facing as the visual field increases in width. Pink traces, Von Mises Mixture model fits of preferred bearing distribution with mean angles depicted and indicated on top left. Bottom row, mean egocentric boundary ratemaps across all EBCs identified for each simulation. Blue to yellow, zero to maximal activity. **b)** Same as in **a**, but for the V1-RSC model.

Furthermore, Figure 7 shows that both models generate more behind-animal EBCs when FOVs are small (60◦, 90° and 120°). Given that there is no mnemonic component in the model and the wall behind the animal is completely out of its view when FOV is small, the result here suggests that the model based on sparse coding promotes the diversity of EBC tuning properties even though only visual input is used.

## 4 Discussion

### 4.1 Summary of key results

In this study, the results of two different learning models for RSC cell responses are compared with experimental RSC cell data. Both models take visual images as the input, using trajectories of the environment that are either measured experimentally or simulated. The Raw Visual (RV) model takes the raw visual images as the input, while the V1-RSC model incorporates visual information processing associated with simple and complex cells of the primary visual cortex (Lian et al., 2019, 2021). After learning, both models generate EBCs that are proximal, distal and inverse, similar to experimentally observed EBCs in the RSC (Alexander et al., 2020). Moreover, the learnt EBCs have similar distributions of orientation and distance coding to the distributions measured in experimental data. The learnt EBCs also show some extent of generalization to novel environments, consistent with the experimental study (Alexander et al., 2020). Furthermore, as the field of view of visual input increases, the orientation distribution of learnt EBCs becomes more lateral. Overall, our results suggest that a simple model based on sparse coding that takes visual input alone can account for the emergence and properties of a special type of spatial cells in the navigational system of the brain - egocentric boundary cells (EBCs). For another recent model that describes the learning of EBCs, see Uria et al.(2022). In the future, this framework can also be used to understand how other visual input (such as landmarks, objects, etc.) affects the firing of spatially-coded neurons, as well as how other sensory input contributes to the tuning properties of some neurons in the navigational system.

### 4.2 Comparison between experimental and model data

Though the model data indicates that both models can learn EBCs similar to experimental ones and the population statistics of orientation and distance coding resembles experimental data, there are still some important differences between model and experimental data that can shed light on the mechanisms associated with EBC responses.

Experimental data shows that the orientation distribution is more skewed towards the back, while the distributions of model data are more lateral (see Figure 5). There are many more behind-animal EBCs in the experimental study compared with the model data when the field of view is 170° (Figure 5, but we found that our model can generate more behind-animal EBCs when the field of view is as narrow as 60° (see Figure 7, suggesting that the competition brought by sparse coding promotes diverse EBC tunings solely based on visual input without any mnemonic component. The difference of population responses among experimental data, RV model data, and V1-RSC model data seems to indicate that a major source of these differences is the extent to which the modelled visual input corresponds to that in the visual system. Whether more biophysically accurate simulated visual input could further reduce these differences is discussed in Section 4.3 & 4.4.

Additionally, there is still a substantial difference in how cells respond in the vicinity of corners of the environment. In simulation, the allocentric ratemaps of some learnt EBCs show overlapping #-like wall responses (see the bottom two examples in Figure 4b and examples in Supplementary Materials A.5), whereas the experimental data seems to “cut off” the segments of #-like response close to the corner. Our models only use visual input while the real animal integrates a variety of different sensory modalities into spatial coding. We infer that the integration of information from different sensory modalities could be responsible for cutting off the overlapping wall responses.

The percentage of EBCs for different data sets also differ, as seen from Table 1. The overall percentage of EBCs was lower in the experimental data than in both types of simulations. This likely arises from the focus of the simulations on coding of static visual input stimuli across a range of different positions and directions in the environment. Though the cells created by this focused simulation show striking similarity to real data, the retrosplenial cortex is clearly involved in additional dimensions of behavior, such as the learning of specific trajectories and associations with specific landmarks. Previous recordings show that neurons in the retrosplenial cortex code additional features such as the position along a trajectory through the environment (Alexander and Nitz, 2015, 2017; Mao et al., 2018, 2020) and the relationship of landmarks to head direction (Jacob et al., 2017; Lozano et al., 2017; Fischer et al., 2020). Human functional imaging also demonstrates coding of position along a trajectory (Chrastil et al., 2015), as well as the relationship of spatial landmarks to specific memories (Epstein et al., 2007). The neuronal populations involved in these additional functions of retrosplenial cortex are not included in the model, which could account for the EBCs making up a larger percentage of the model neurons in the simulations.

### 4.3 Rat vision processing

Rats have very different vision from humans, in part because their eyes are positioned on the side of their head, whereas human’s eyes are facing front. Consequently rats have a wide visual field and a strong lateral vision. In this study, rat vision is simulated by a camera with a 170° horizontal view and 110° vertical view, except for the results in Section 3.3. In Section 3.3, when different horizontal fields of view are used, we found that the model can generate more behind-animal EBCs with smaller field of view and the orientation becomes more lateralized as the field of view increases. Though the view angle of 170° is wider compared with human vision, the simulated vision might not be as lateral as in real rats. Due to the built-in limitations of the Panda3D game engine used to simulate the visual input, we were unable to generate visual input at degrees more lateral than the 170 degree range used here. Additionally, real rats have binocular vision instead of a monocular vision simulated in this study. This will be investigated in future studies, in which the rat vision will be mimicked by simulating visual input using two laterally positioned cameras. As a more biophysically accurate simulated visual input is used, we infer that this could further reduce differences between model and experimental data, including generating more behind-animal EBCs when the field of view is large.

### 4.4 Differences between Raw Visual model and V1-RSC model

Both the RV and V1-RSC models take the visual input and generate EBC responses using learning methods based on the principle of sparse coding. However, there are significant differences between the two models. The RV model takes the raw image as the input while the V1-RSC model incorporates vision processing similar to that of the brain that detects lines or edges in the visual input. In other words, the RV model learns cells based on the individual pixel intensities while the V1-RSC model learns cells based on the existence of visual features such as lines or edges. Because the environment consists of three black walls and one white wall, this difference may result in the white wall affecting the RV model more than the V1-RSC model. In particular, this could explain why the learnt EBCs of the V1-RSC model tend to be more omnidirectional in their firing for all four walls compared with the RV model (see examples of both models in Supplementary Materials), which may be related to the role of RSC as the egocentric-allocentric “transformation circuit” proposed by Byrne et al.(2007) and Bicanski and Burgess (2018) that transforms upstream egocentric sensory responses (vision in this paper) into downstream allocentric spatial cells. Another difference lies in the percentage of learnt EBCs between two models, where the V1-RSC model learns more EBCs (see “Comparison between experimental and model data”, Section 4.2, above). We infer that this difference also originates from the different visual input processing carried out in the models. Geometries (lines or edges) seem to be important for the EBCs firing, so the ability to detect such features in the V1-RSC model may help the model learn more EBCs. In addition, the RV model shows more diverse tuning properties of learnt EBC population than the V1-RSC model (see Figure 5), while the V1-RSC model shows better generalization to novel environments (see Figure 6), likely caused by the V1 pre-processing of the model. Differences between the responses in the two models also point to the effect that the processing of visual input carried out in the early visual pathway (retina to primary visual cortex) has upon RSC cell responses (Lian et al., 2019, 2021). Since the V1-RSC model is a better model of rat’s vision processing system, we infer that its model EBCs will be more similar to EBCs in the brain (also see Section 4.3, above). Furthermore, the model will better account for experimental data as a more biophysically accurate simulated visual input is used.

## Author Contribution Statement

YL, SW, ASA, MEH and ANB conceived the work. YL and SW designed the model. ASA and MEH provided experimental data. YL, SW and ASA analysed the model results. YL and SW wrote the first draft of the manuscript. All authors participated in writing and editing of the manuscript.

## Acknowledgements

This work received funding from the Australian Government, via grant AUS-MURIB000001 associated with ONR MURI grant N00014-19-1-2571. This research was also supported by NIH NINDS K99 NS119665, NIMH R01 MH120073; Office of Naval Research MURI grant N00014-16-1-2832; Office of Naval Research MURI N00014-19-1-2571; and Office of Naval Research DURIP N00014-17-1-2304.

## Code Availability

The code of implementing the model is made available at https://github.com/yanbolian/ Learning-EBCs-from-Visual-Input.

## A Supplementary Materials

A.1–A.4 of this section provide all learnt cells of both Models recovered by simulated and experimental trajectories.A.5 provides two examples of learnt EBCs of V1-RSC model that show overlapping wall response in their ratemaps. Each row with three images below and in the following subsections shows the spatial ratemap, firing plot with head directions and egocentric ratemap.

### A.1 All learnt cells of Raw Visual (RV) model using experimental trajectory

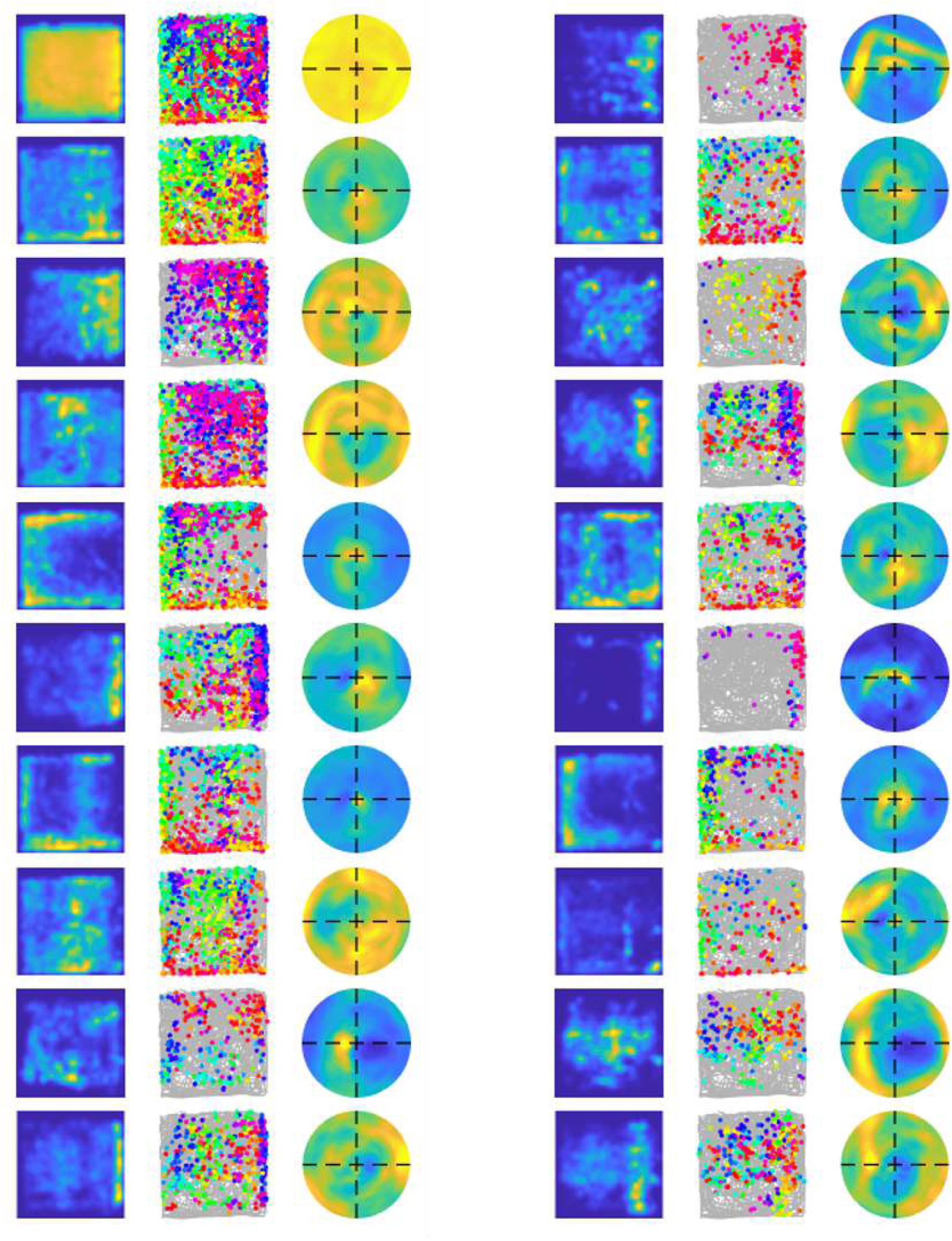
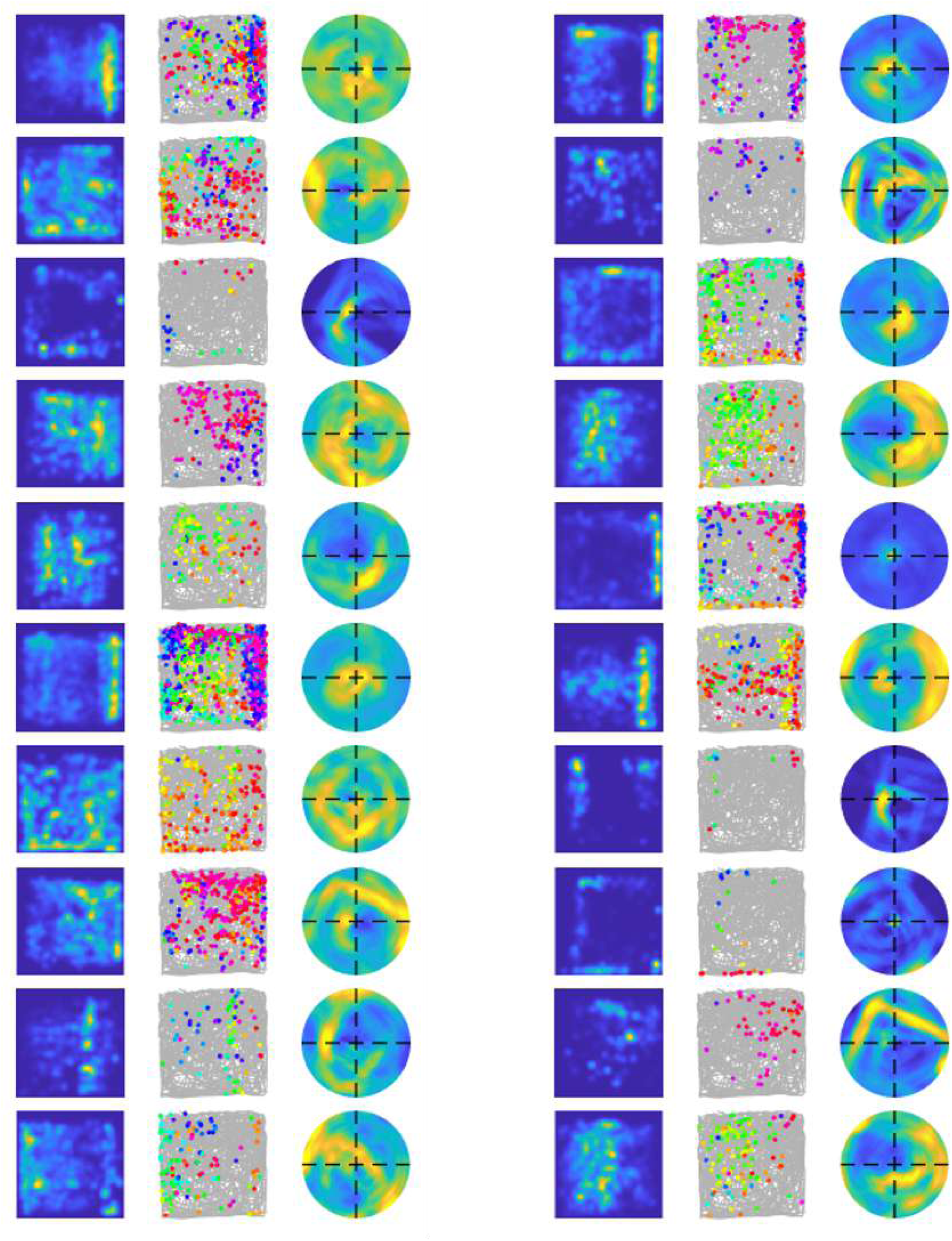
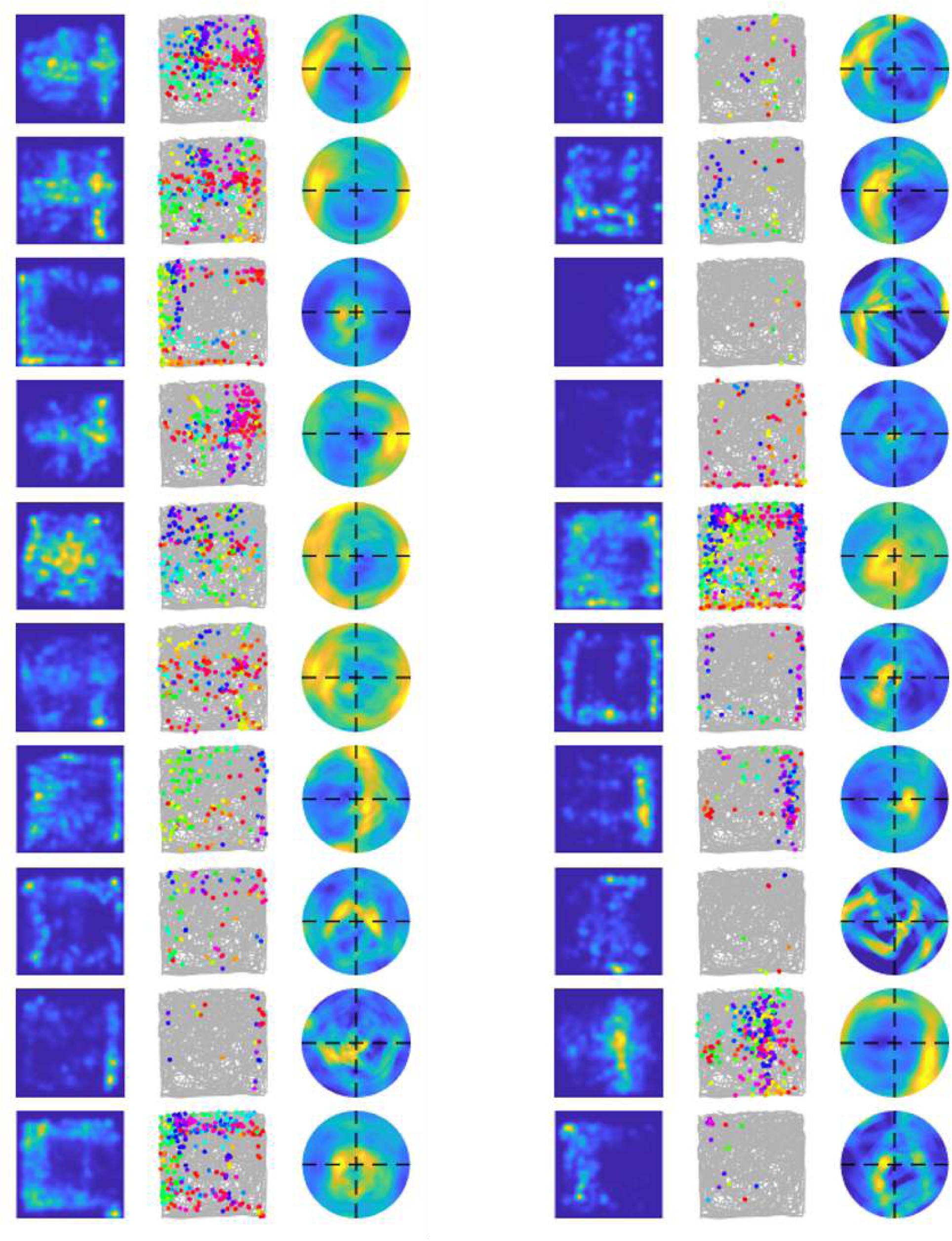
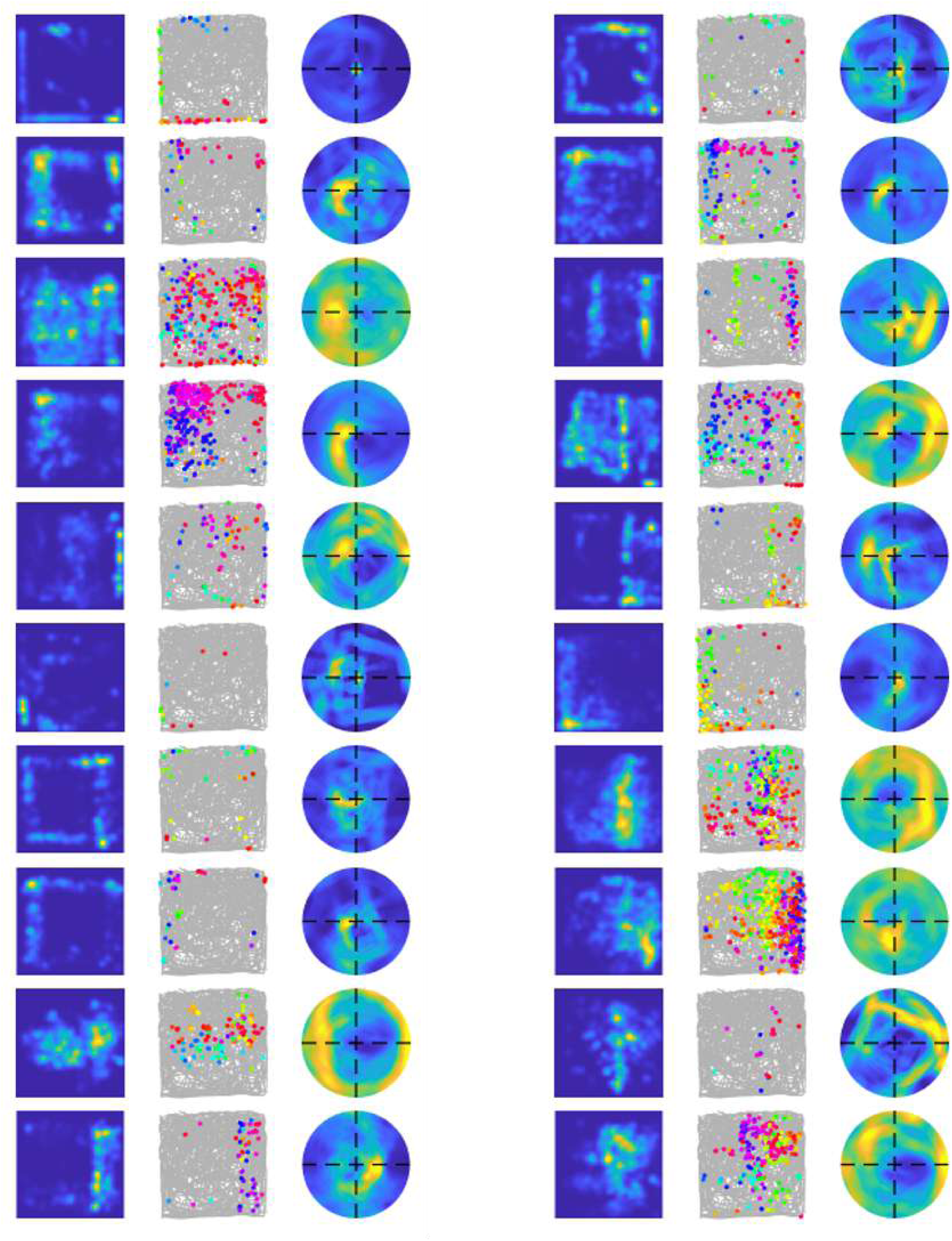
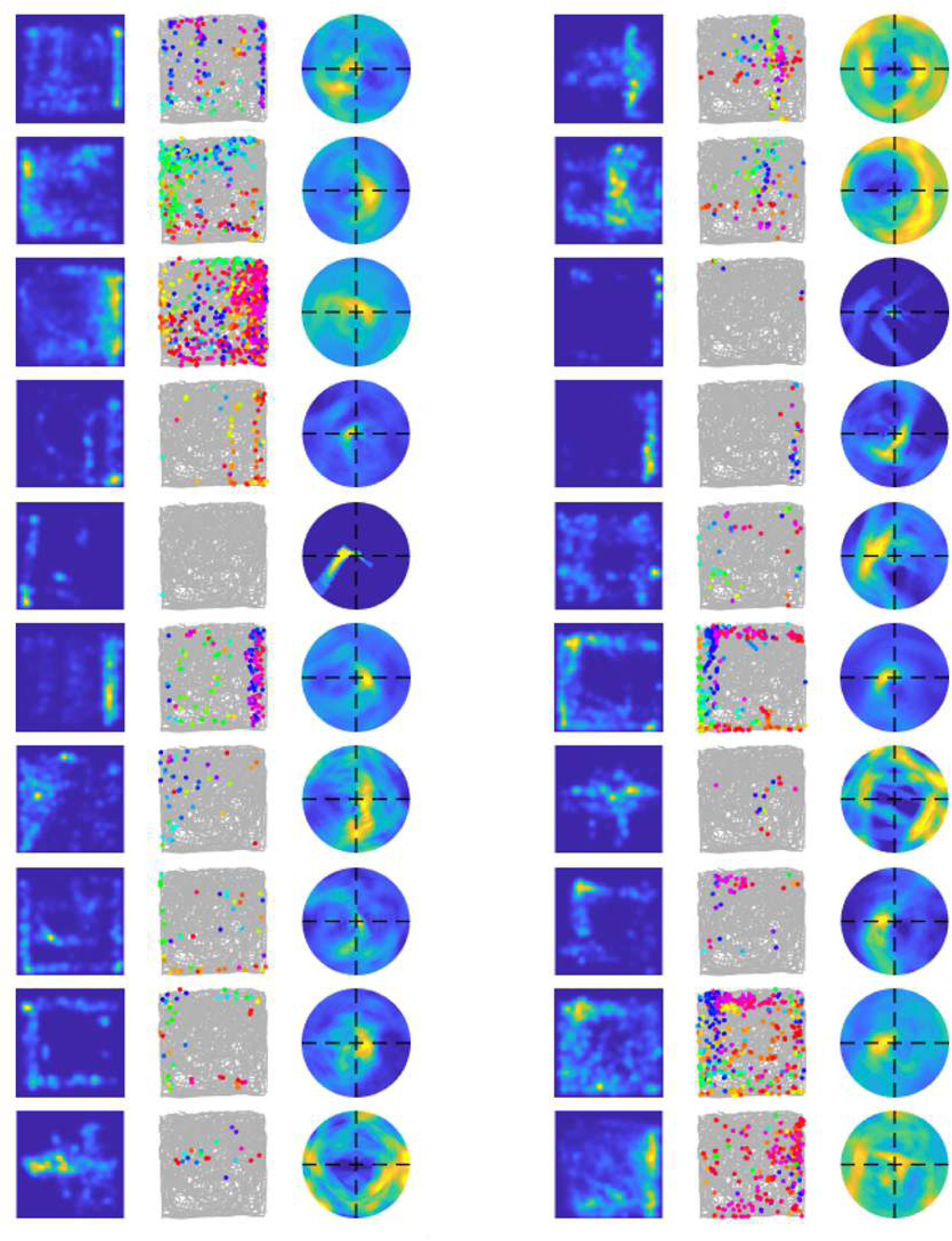

### A.2 All learnt cells of Raw Visual (RV) model using simulated trajectory

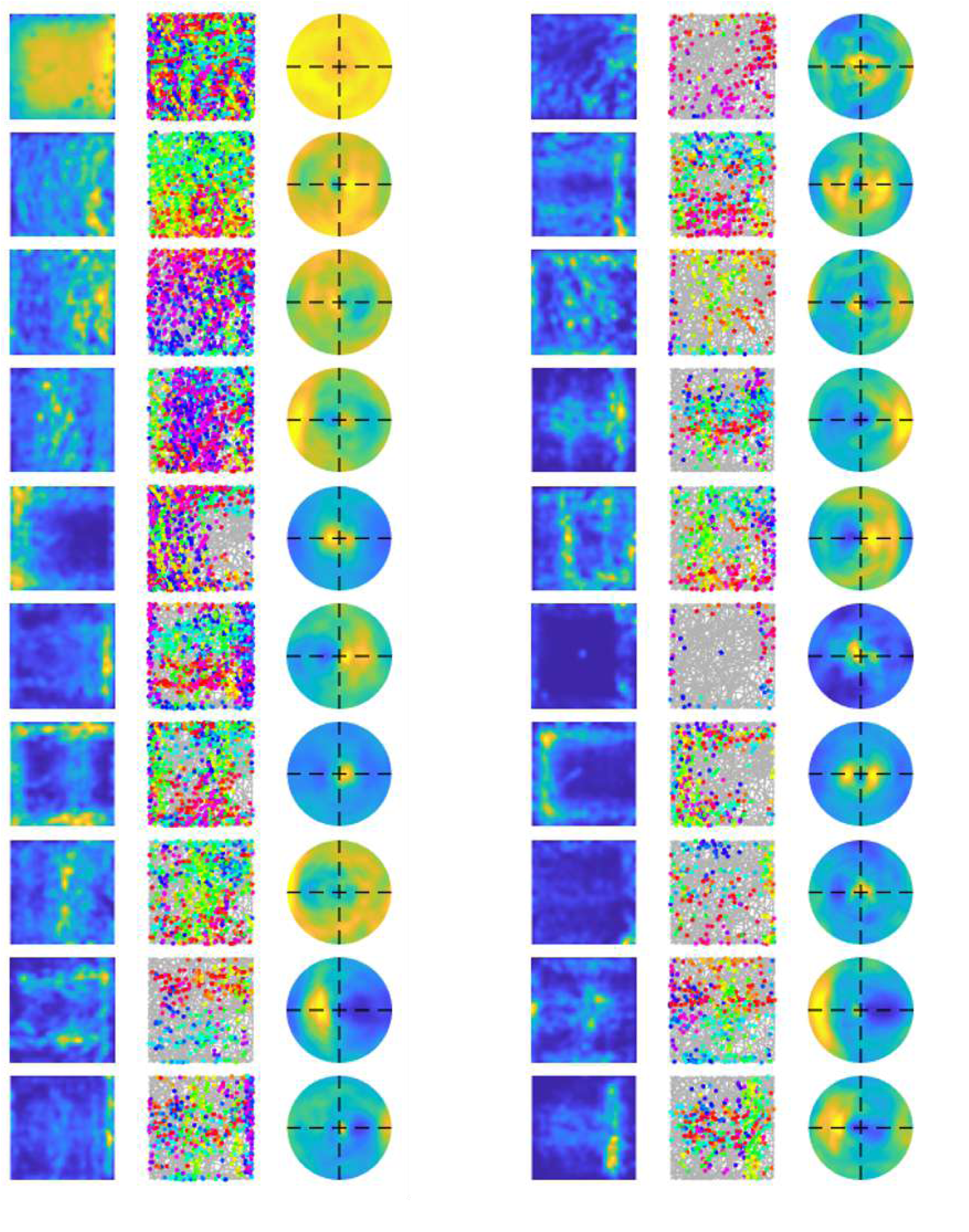
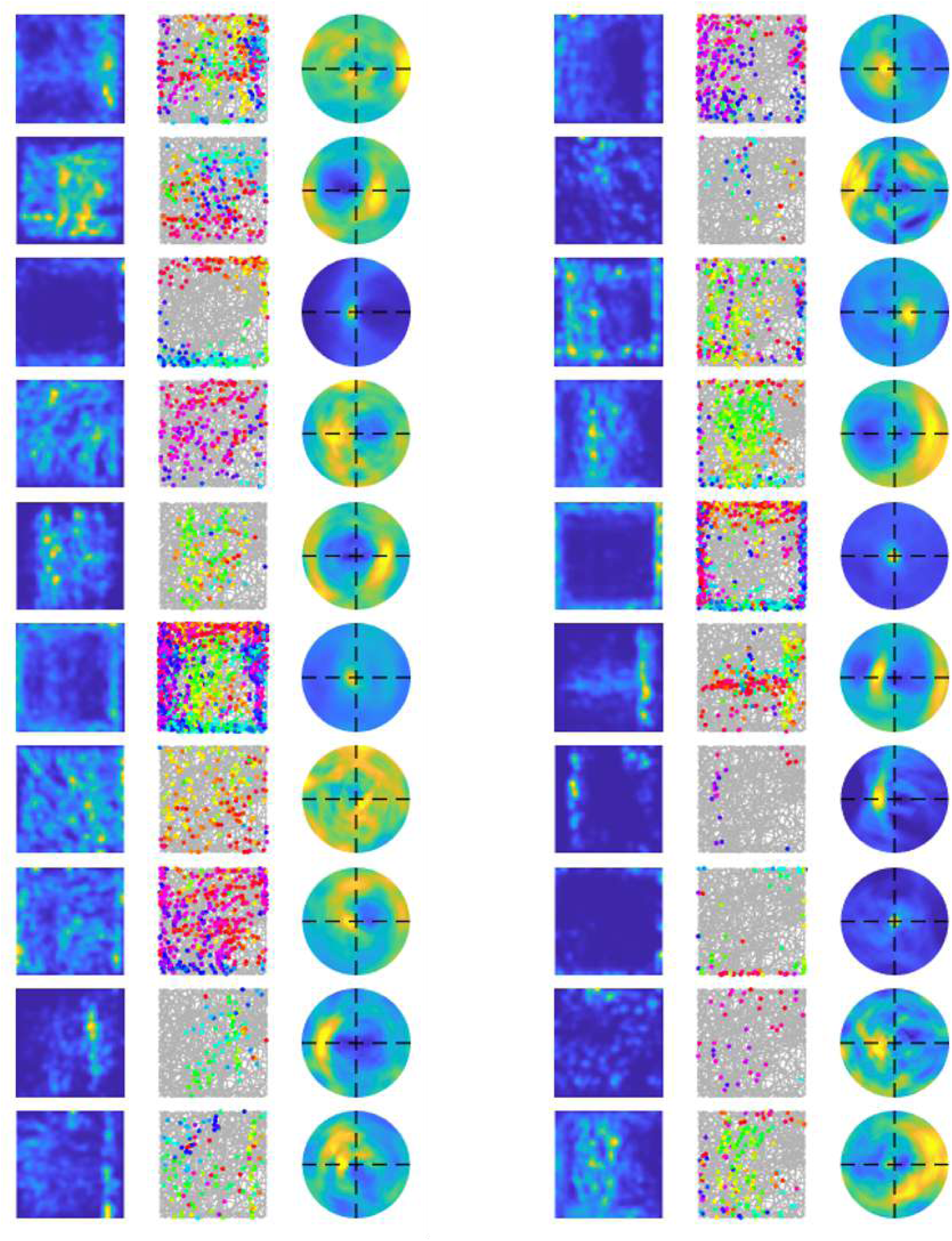
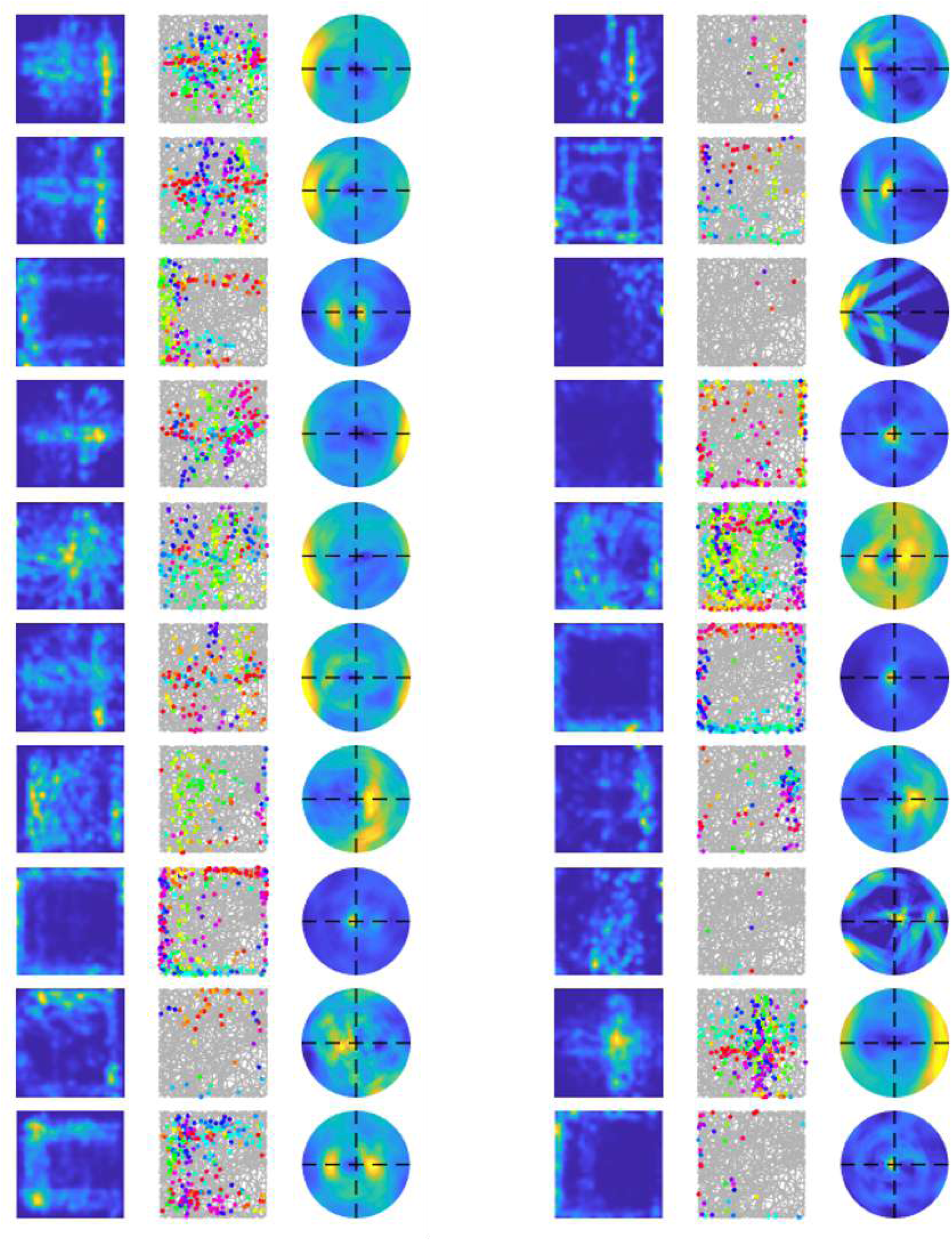
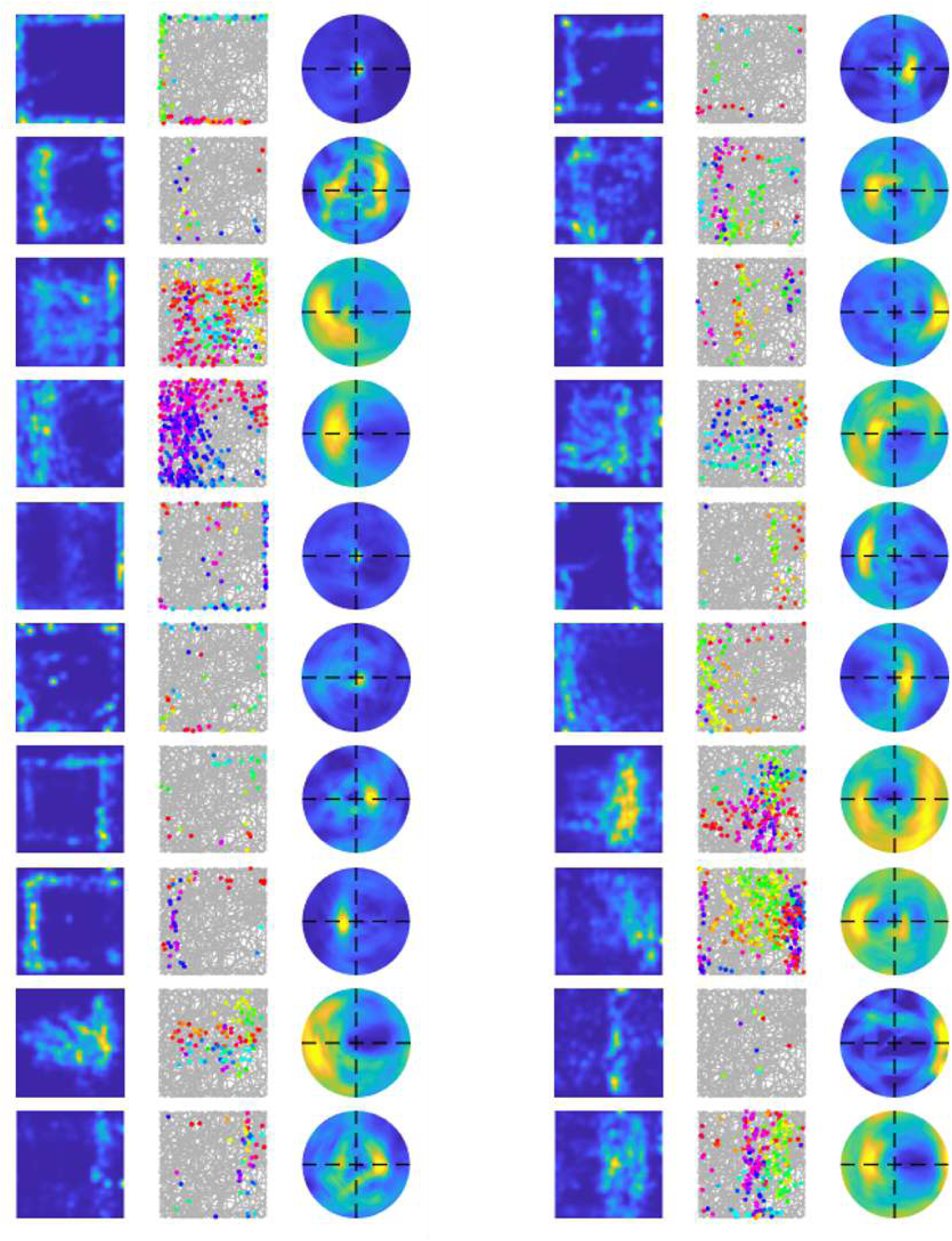
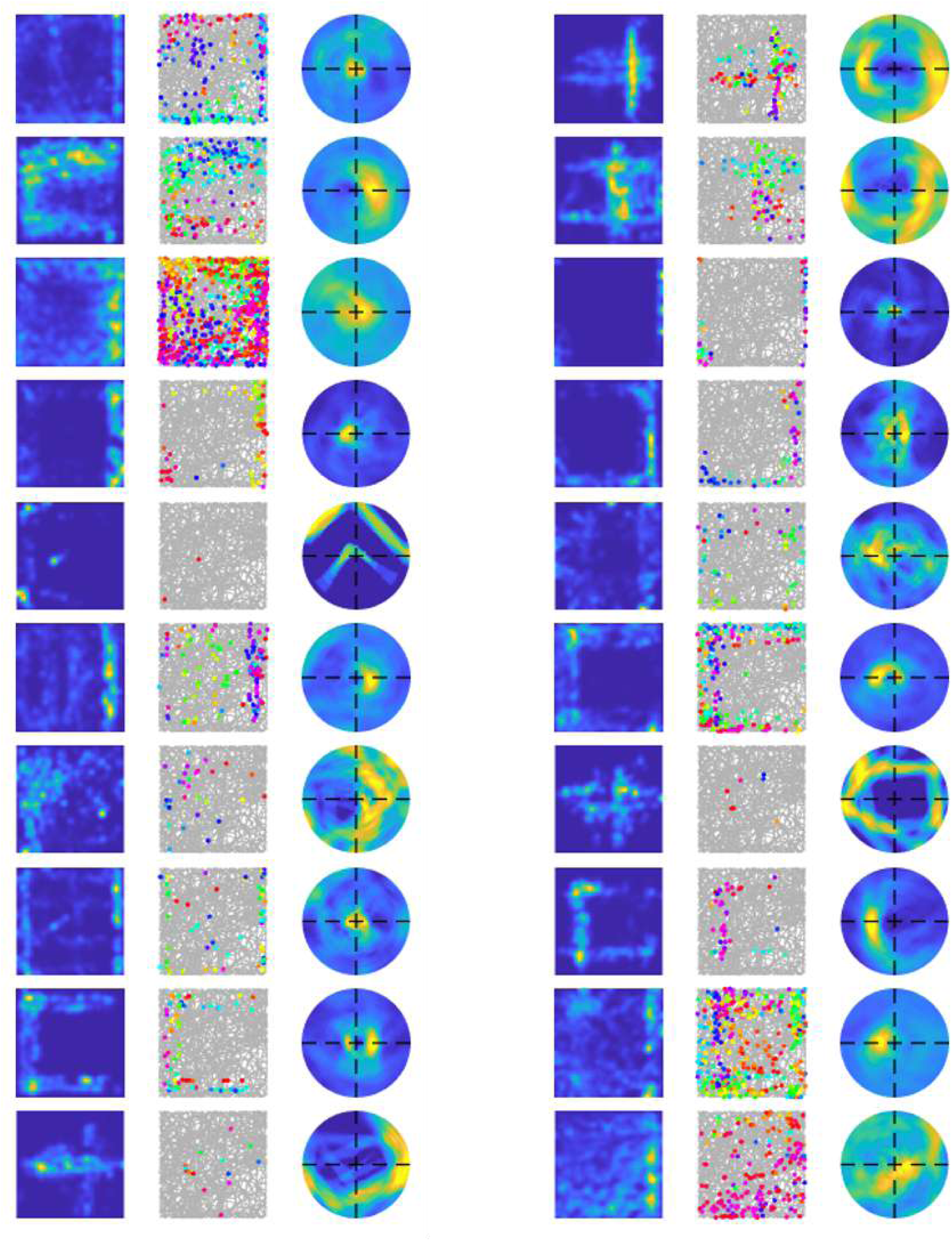

### A.3 All learnt cells of V1-RSC model using experimental trajectory

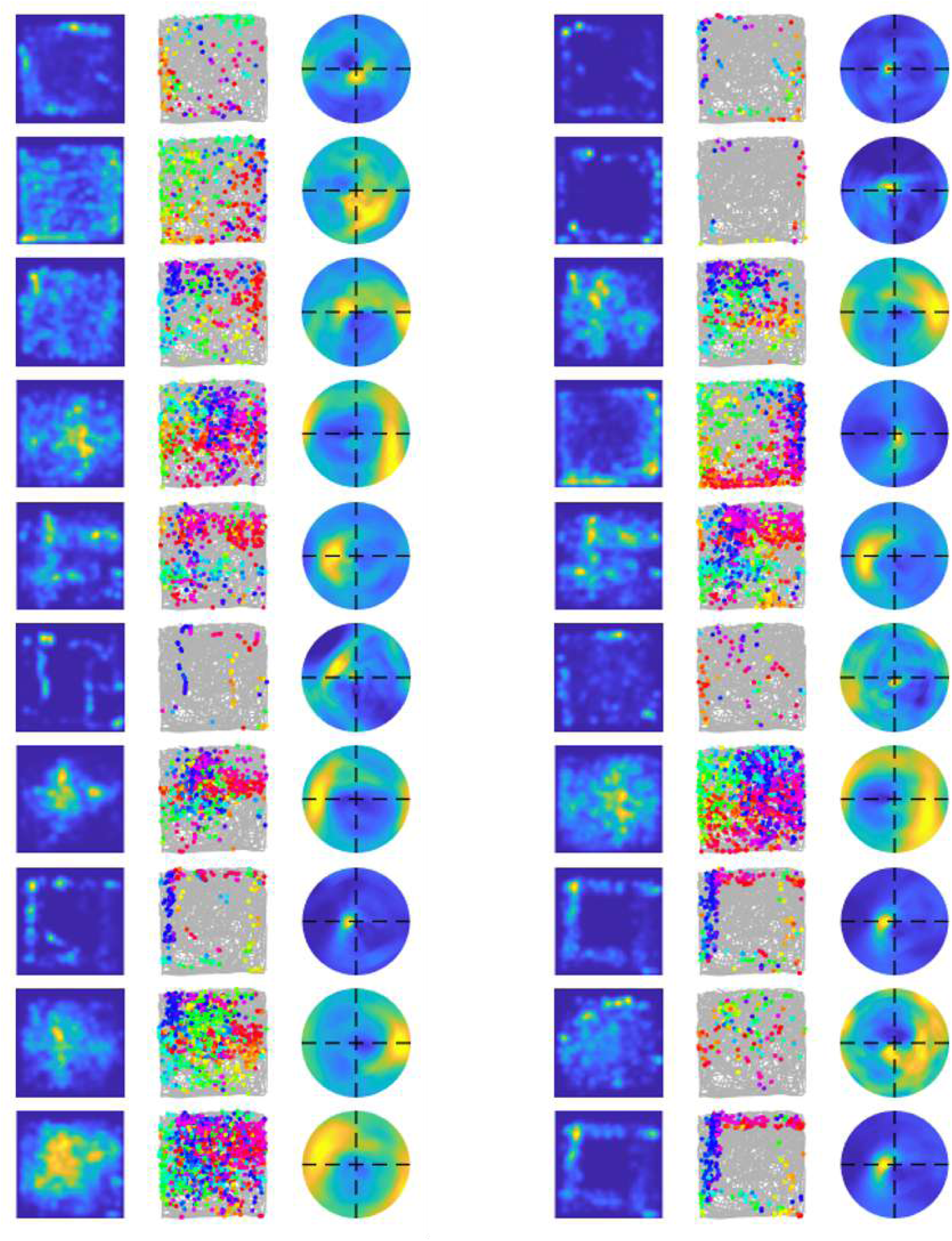
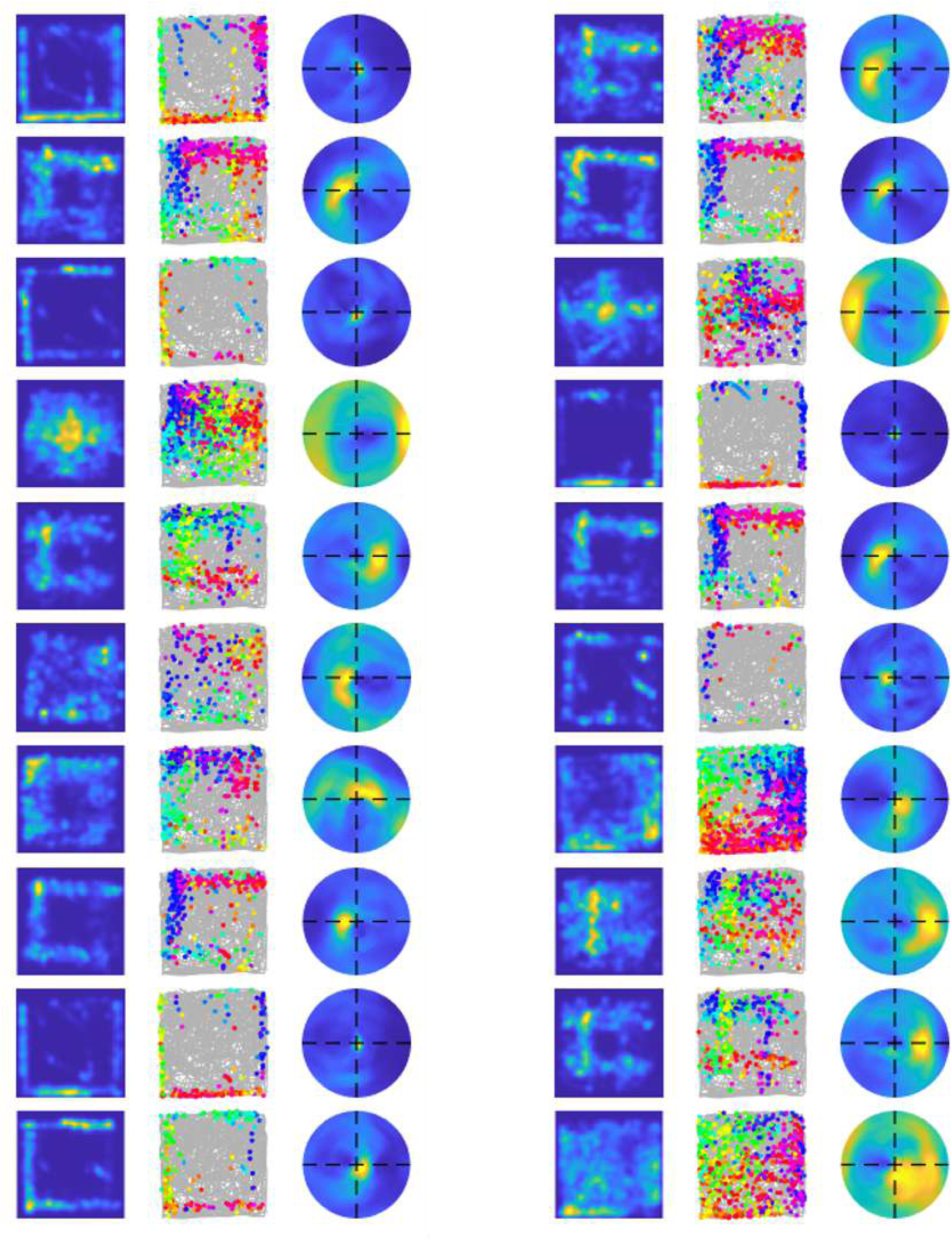
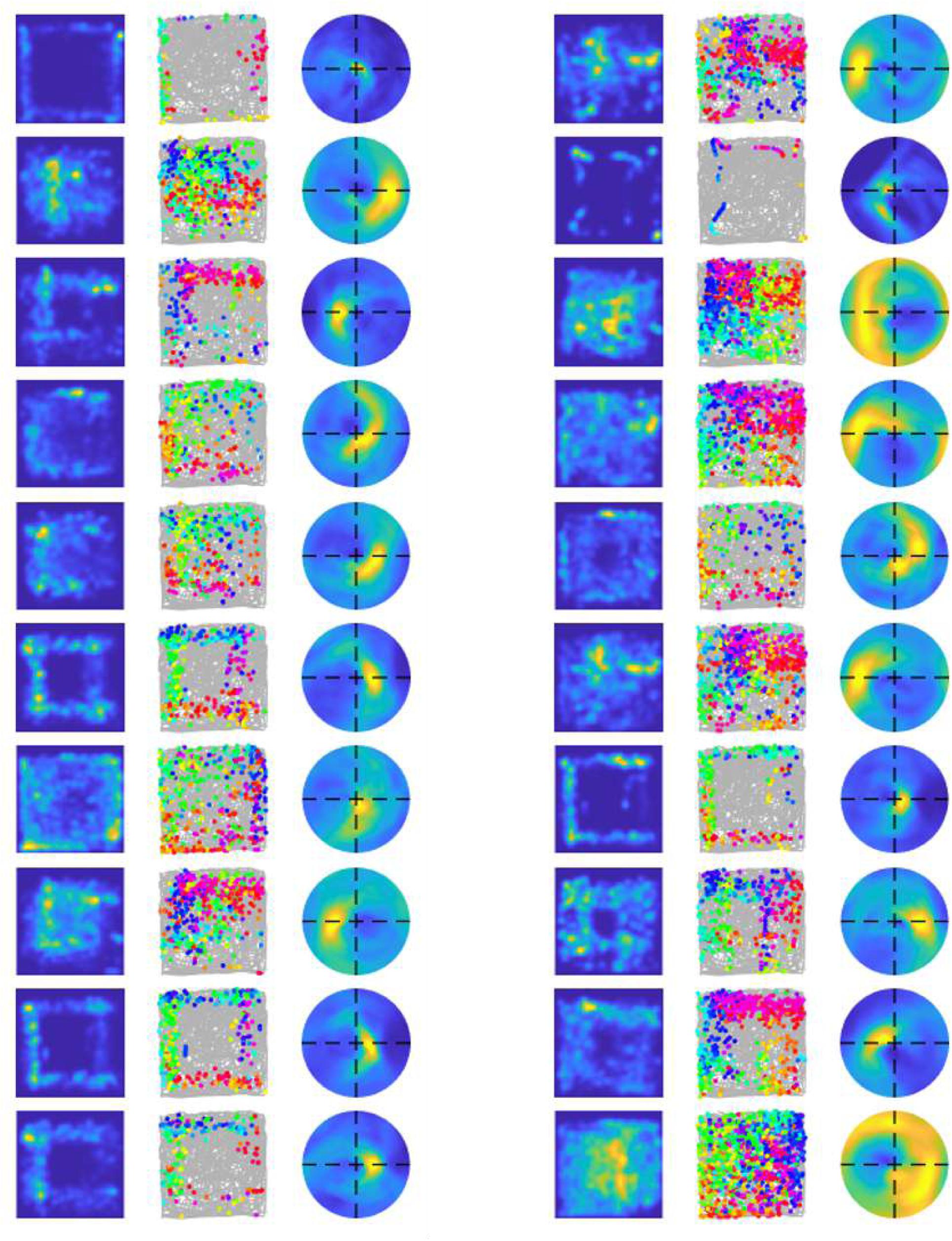
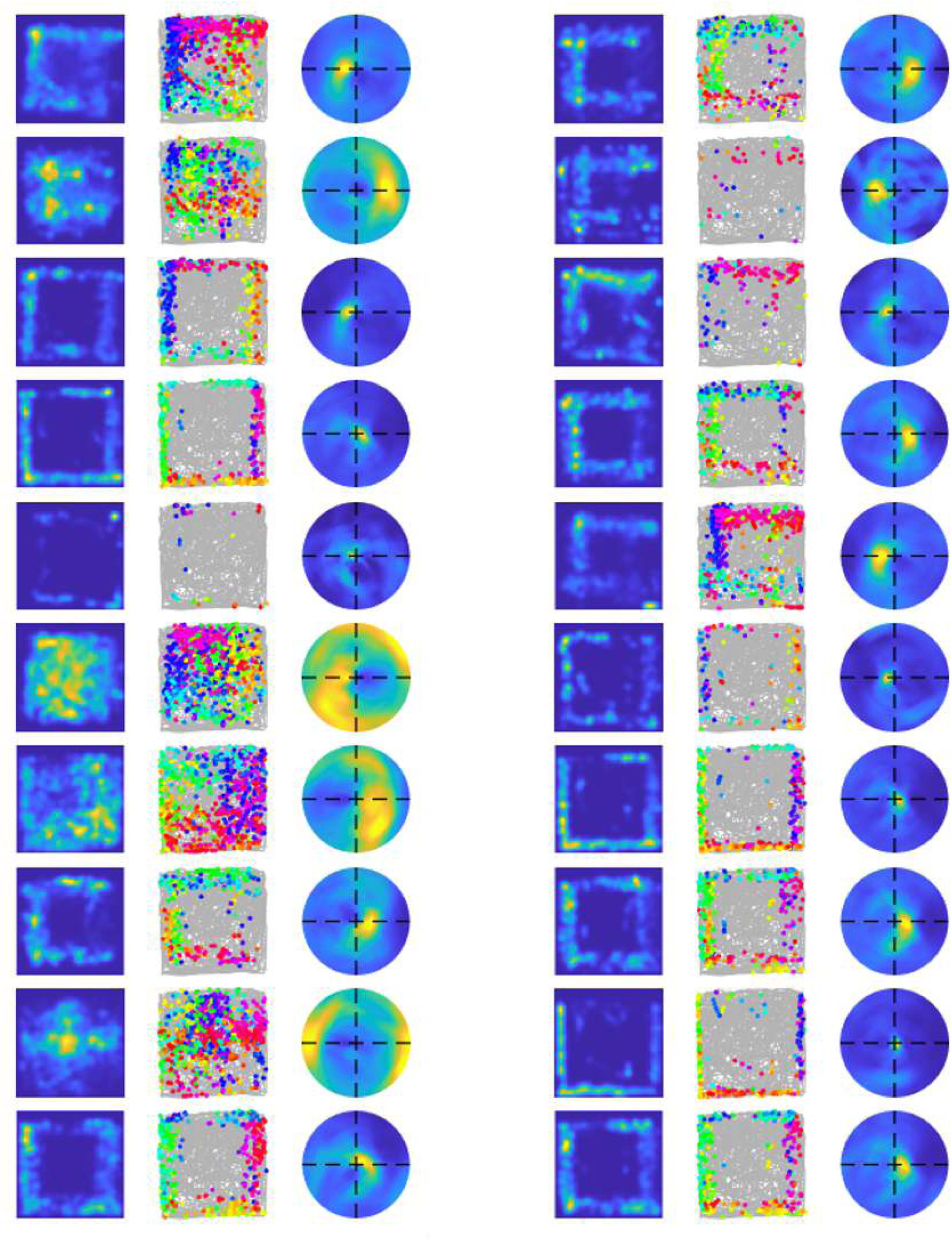
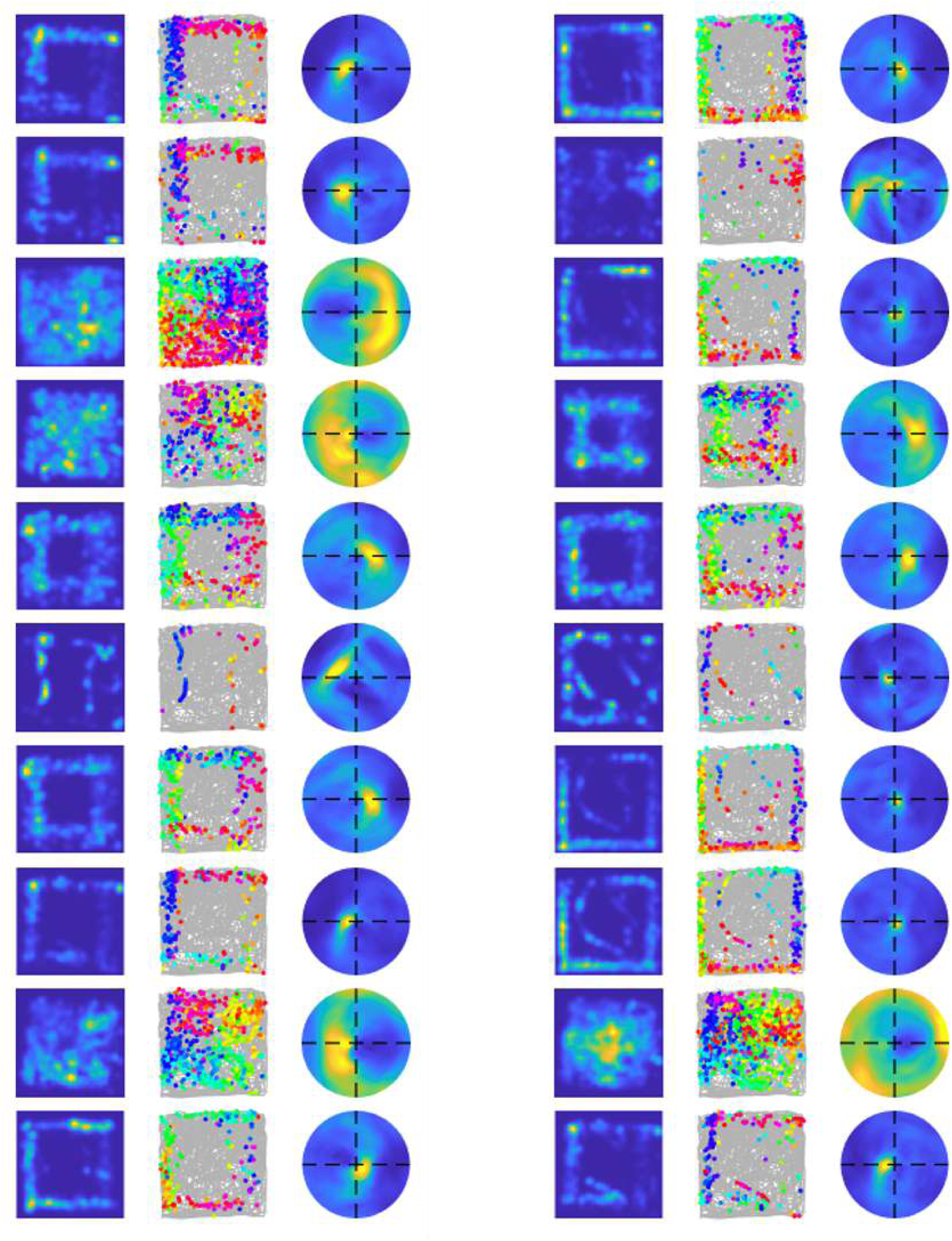

### A.4 All learnt cells of V1-RSC model using simulated trajectory

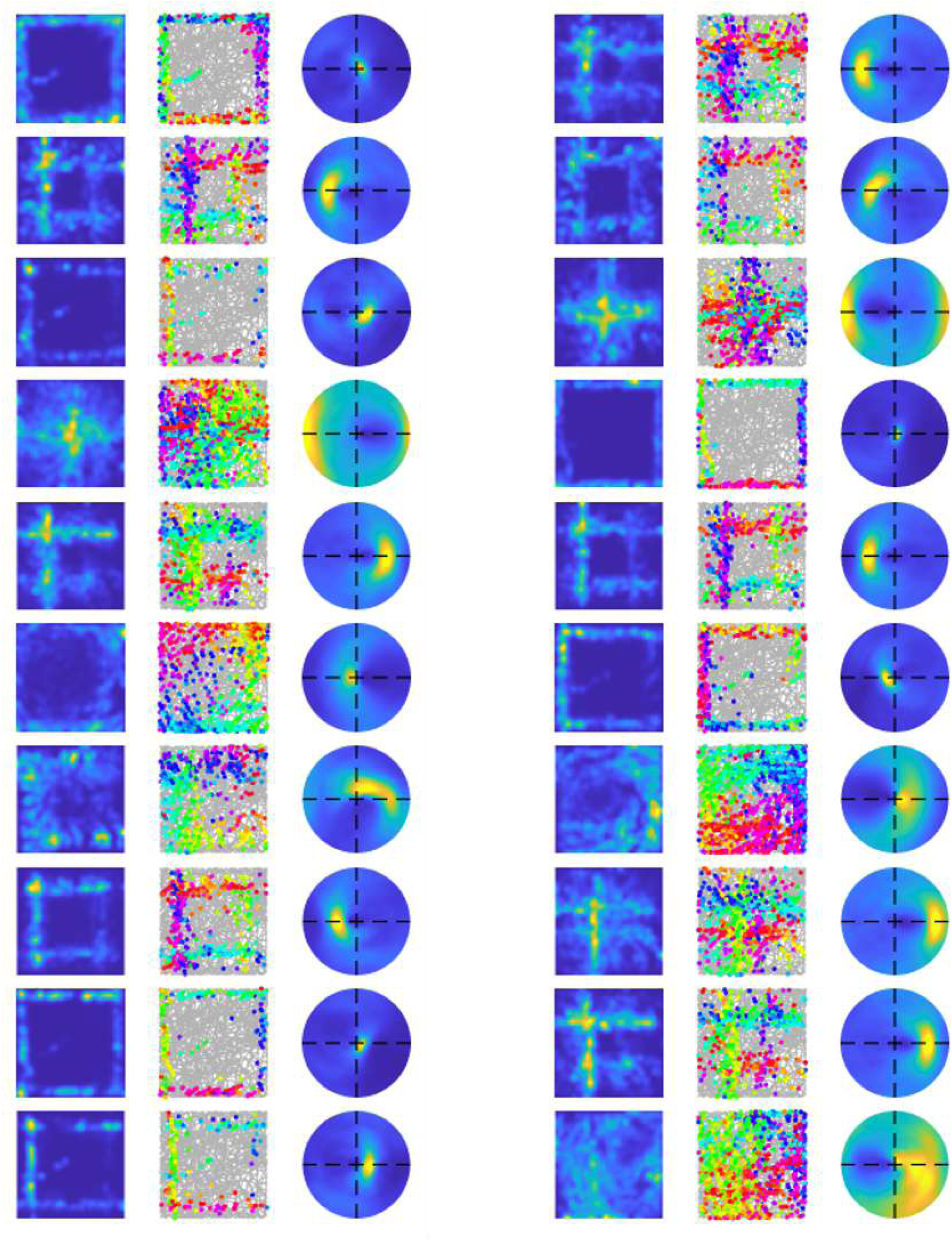
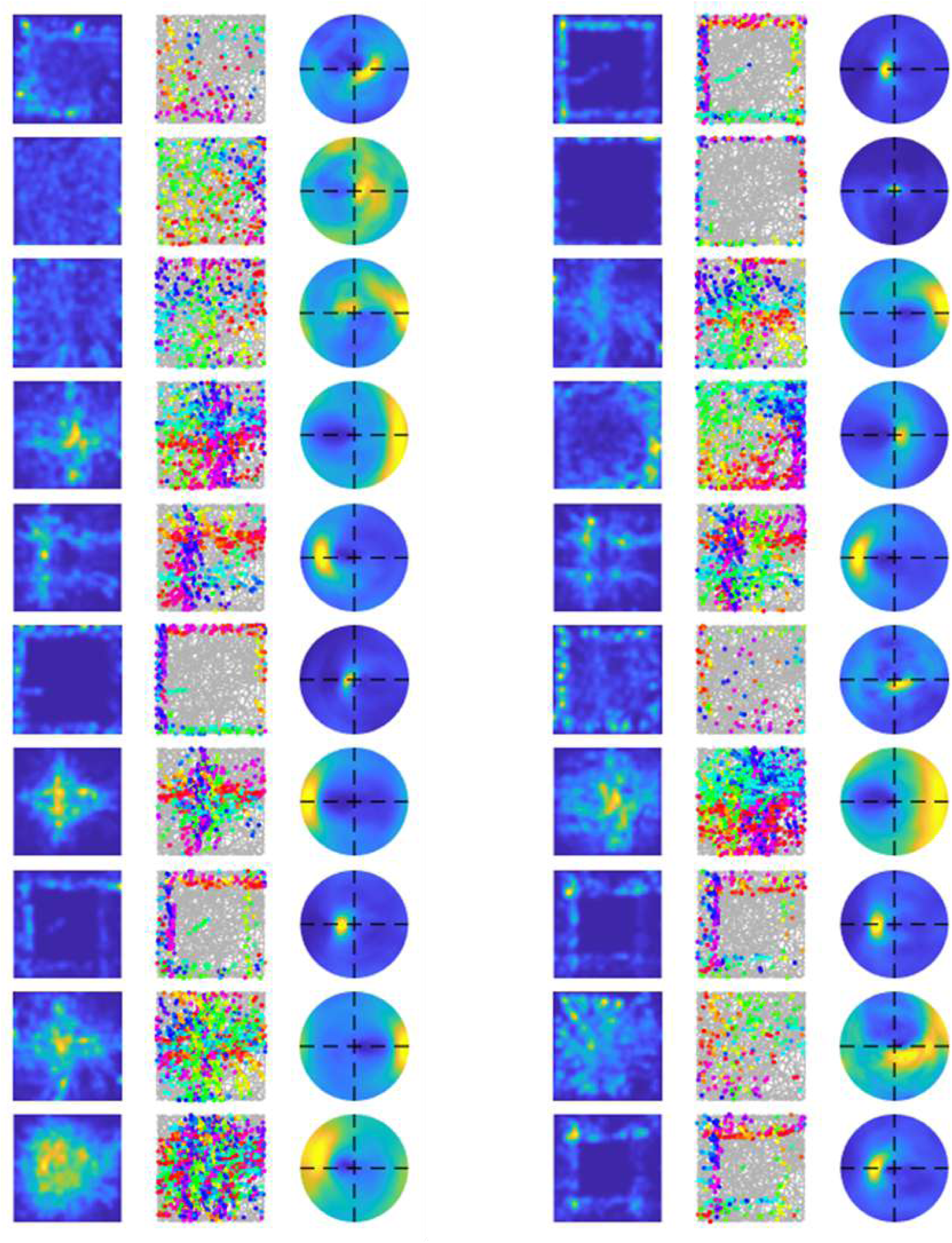
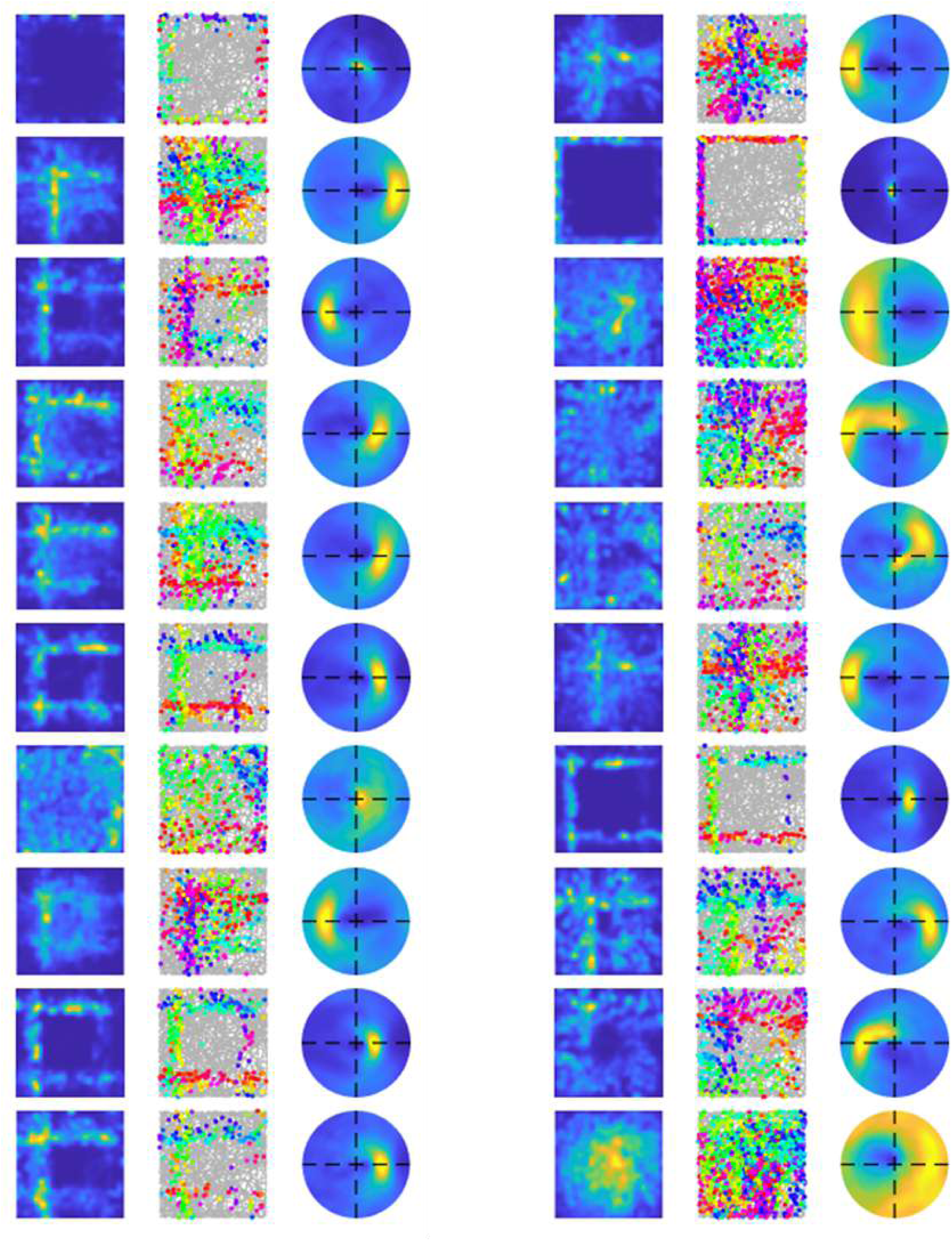
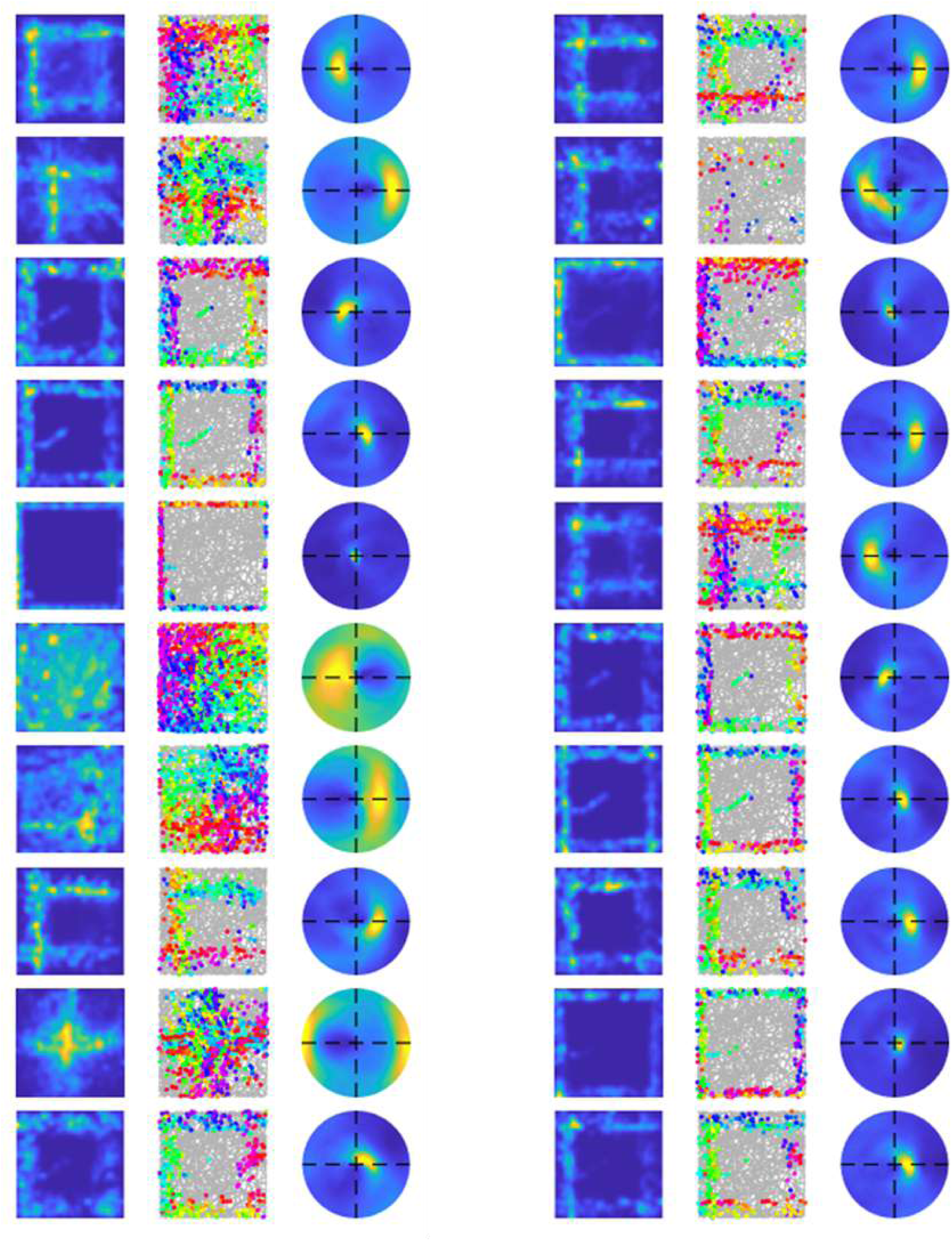
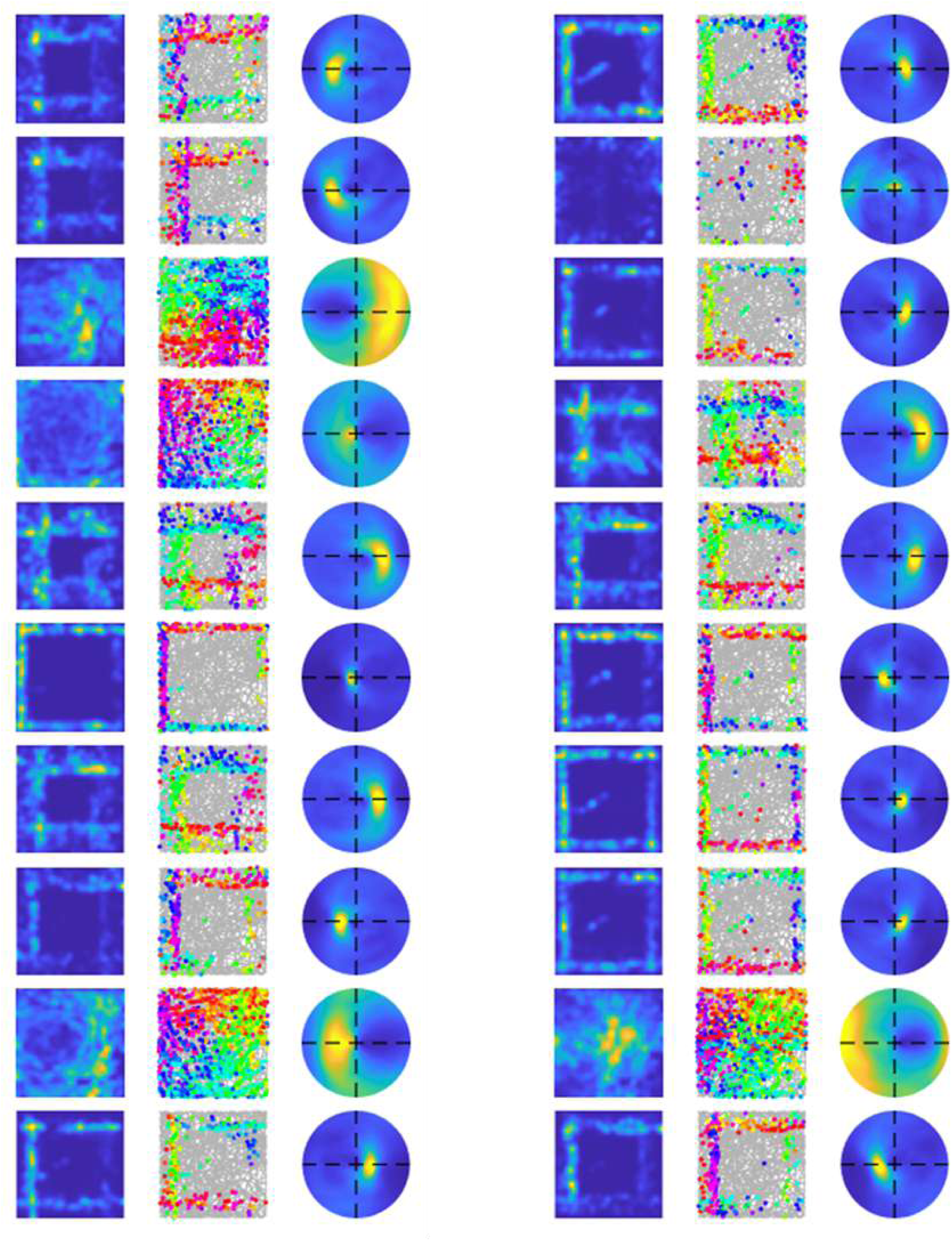

### A.5 Examples of learnt EBCs that show overlapping wall response

Plotted here are two examples of learnt EBCs that show overlapping wall response in their ratemaps. Each row with three images shows the spatial ratemap, firing plot with head directions, and egocentric ratemap. These two examples of learnt EBCs from the V1-RSC model do not “cut off” the segments close to the corner such that the spatial ratemaps have overlapping #-like responses.

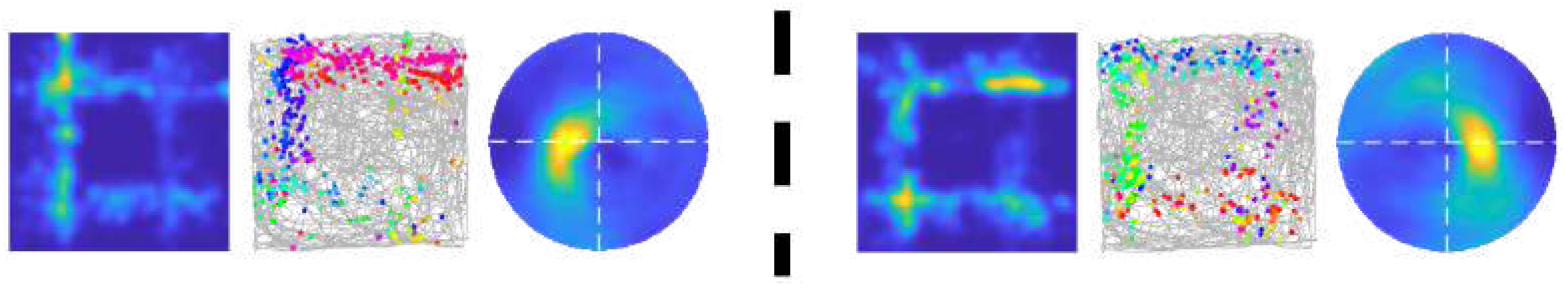

